# Structural insights into inhibition mechanism of the helicase-primase complex from human herpesvirus 1

**DOI:** 10.1101/2025.05.12.653380

**Authors:** Ko Sato, Hisashi Ishida, Toma Miyagishi, Shunsuke Kobayashi, Yoshiaki Kise, Keisuke Hamada, Chikako Okada, Asako Oguni, Osamu Nureki, Hidetoshi Kono, Kaori Fukuzawa, Toru Sengoku

## Abstract

Human herpesviruses (HHVs) are widespread pathogens causing severe diseases, especially in immunocompromised individuals, and are linked to neurodegenerative and autoimmune disorders. Current antiviral therapies primarily target the α-subfamily, with limited options for β- and γ-subfamilies. The helicase-primase complex (HPC) is essential for viral DNA replication and represents a key antiviral target, but its structure and inhibition mechanisms were previously unclear. Here, cryo-EM structures of the HHV1 (α-subfamily) HPC bound to single-stranded DNA and two clinical inhibitors, amenamevir and pritelivir, are reported. The HPC features flexible helicase and primase modules, with both inhibitors binding a shared allosteric pocket in the helicase module. Structural, biochemical, and molecular dynamics analyses indicate that these inhibitors lock the helicase in an open, inactive conformation. Combined molecular dynamics and fragment molecular orbital analyses explain the α-subfamily selectivity and distinct antiviral spectra of these drugs. These findings provide a structural framework to guide the development of novel inhibitors targeting β- and γ-subfamily herpesviruses.

**Significance:** Herpesviruses are ubiquitous human pathogens responsible for a wide spectrum of diseases, including severe infections in immunocompromised individuals and associations with neurodegenerative and autoimmune disorders. Despite the clinical importance of these viruses, current antiviral therapies are largely limited to the α-subfamily, with few effective options for β- and γ-subfamilies. The helicase-primase complex (HPC) is a critical enzyme for viral DNA replication and a validated antiviral target, yet its structural basis and inhibition mechanisms have remained elusive, hindering the rational development of next-generation therapeutics. This study provides the first high-resolution cryo-EM structures of the human herpesvirus 1 (HHV1) HPC bound to single-stranded DNA and two clinically relevant inhibitors, amenamevir and pritelivir. These structures reveal the modular organization of the HPC and identify a shared allosteric pocket where both inhibitors bind. Structural, biochemical, and molecular dynamics analyses demonstrate that these inhibitors lock the helicase in an open, catalytically inactive conformation, thereby blocking ATP binding and halting DNA unwinding. Fragment molecular orbital (FMO) calculations further elucidate the molecular determinants of inhibitor binding, explaining the α-subfamily specificity and the distinct antiviral spectra of amenamevir and pritelivir. The integrated approach combining cryo-EM, molecular dynamics, and quantum chemical calculations not only clarifies the mechanism of action of current clinical inhibitors but also establishes a robust framework for the rational design of novel antivirals. By providing detailed insights into the structural and dynamic properties of the HPC and its inhibition, this work paves the way for the development of new drugs with tailored specificity and improved pharmacokinetic properties, potentially expanding therapeutic options to currently untreatable herpesvirus infections. These findings have broad implications for antiviral drug discovery and for understanding the molecular mechanisms underlying DNA replication in herpesviruses.

## Introduction

Human Herpesviruses (HHVs) are significant human pathogens^1^. To date, eight herpesviruses (HHV1–8) have been identified and classified into three subfamilies: α, β, and γ. All these herpesviruses can cause human diseases, including herpes simplex, herpes zoster, encephalitis, and various tumors. They can also lead to severe conditions, particularly in immunocompromised individuals. Moreover, recent studies have suggested potential links between herpesvirus infections and chronic diseases such as Alzheimer’s disease^2^, dementia^3^, multiple sclerosis^4^, and chronic fatigue syndrome^5^. Currently, multiple anti-herpes drugs, such as nucleoside analogs and helicase-primase inhibitors (HPIs), are available for the α-subfamily (HHV1/HSV1, HHV2/HSV2, and HHV3/VZV). In contrast, therapeutic options for the β-subfamily (HHV4/Epstein-Barr virus and HHV8/Kaposi’s sarcoma-associated virus) and γ-subfamily (HHV5/cytomegalovirus, HHV6, and HHV7) remain limited, underscoring the need to develop new antiviral drugs targeting these subfamilies.

Herpesviruses employ a set of unique enzymes for the replication of their genomes, which can be therapeutically targeted by specific inhibitors. One such target is the helicase-primase complex (HPC)^6^, composed of three subunits: the helicase subunit (UL5 in HHV1), the primase subunit (UL52 in HHV1), and a non-catalytic accessory subunit (UL8 in HHV1). The HPC associates with the replication fork and unwinds the DNA duplex through the action of the helicase subunit, generating ssDNA as the templates for DNA synthesis by the viral DNA polymerase. While most replicative helicases in eukaryotes and prokaryotes form hexameric ring structures surrounding the DNA^7^, herpesvirus helicases belong to superfamily Ib, which is thought to function as monomers^8^. A previous studies on a related superfamily Ib helicase has shown that they alternate between “open” and “closed” conformations in an ATP hydrolysis-dependent manner, translocating along ssDNA in an inchworm-like motion to separate the base pairs they encounter^9^. The HPC also synthesizes an RNA primer on the lagging strand through the action of the primase subunit. This subunit contains sequence motifs conserved among archaeo-eukaryotic primases^10^, along with a putative zinc-binding domain (ZBD), which is commonly found in other primases, including the human PrimPol protein. Notably, all three subunits of the HPC are required for the optimal activities of both the helicase and primase, suggesting complex regulatory interactions between them. As no experimental structure of the HPC has been determined, it remains elusive how these subunits interact with each other, how the two catalytic domains are spatially arranged, and how the two activities are coordinated within this unique complex.

HPIs specifically inhibit the HPC of α-subfamily, thereby suppressing their replication. One such molecule, amenamevir (AMNV)^11,12^ has demonstrated efficacy against all three α-herpesviruses, HHV1 and 2 (also known as herpes simplex viruses 1 and 2, respectively), and HHV3 (varicella zoster virus). AMNV has been approved in Japan for the treatment of herpes simplex and herpes zoster; however, its poor brain penetration^13^ limits its clinical utility for treating herpesvirus-induced encephalitis. Pritelivir (PTV)^14,15^, another clinical HPI in phase 3, is effective against HHV1 and HHV2, but shows only limited activity against HHV3 and HHV5^14^. To date, no HPIs effective against β- and γ-subfamily herpesviruses have been reported. The lack of structural analysis of the interaction between α-herpesvirus HPC and HPI has hindered rational design of next-generation HPIs effective against β- and γ-subfamily herpesviruses.

## Results

### Overall structure

To elucidate the overall architecture of the HPC and the inhibition mechanism by HPIs, we set out to determine its cryo-EM structures. We reconstituted the HPC of HHV1 by co-expressing the three subunits in insect cells and purified the complex by affinity chromatography. We mixed the purified HPC with a 43-mer ssDNA, AMP-PNP (a nonhydrolyzable analog of ATP) and either AMNV or PTV, and collected cryo-EM images. During the EM volume reconstruction, we noticed that the HPC can be divided into two modules, hereafter referred to as the helicase module (HM), consisting of the entire UL5 (helicase) and two domains of UL52 (primase), and the primase module (PM), consisting of the primase domain of UL52 and the entire UL8 (non-catalytic subunit). The relative positions of the two modules are flexible. Thus, following the global density reconstruction, focused refinement of the two modules was independently performed, yielding two locally refined maps with improved resolution (about 2.9-3.3 Å), on which we built atomic models (Figure S1-S5). We also performed a 3DFlex refinement^16^ of the overall complex which gave a better overall resolution (about 3.1-3.3 Å) and built the full atomic models into these 3DFlex maps with the help of the two atomic models built on locally refined maps (Supplementary Movie 1). Unless otherwise stated, we use locally built models to describe local interactions and the 3DFlex-derived models for discussion on global structures.

Figure 1 and Figure S6 show the overall structures of the AMNV- and PTV-bound complexes, respectively. The HM consists of the entire UL5 protein and two domains of UL52, namely the scaffold domain (SD, residues 1-408 and 891-910) and the ZBD (residues 911-1043). The two domains of UL52 intimately interact with UL5 with large interface areas. UL5 consists of the domains 1A (residues 34-344 and 864-892), 2A (residues 345-417 and 805-863), 2B (residues 418-514 and 693-804), and 2C (residues 515-692), which are shared by superfamily Ib helicases. Structure similarity analysis using the DALI server^17^ revealed that the helicase subunit is most similar to Pif1 helicases (e.g., PDB ID: 7OTJ; Z score = 19.2)^18–21^. Of the 43-mer ssDNA used for structural determination, only seven nucleotides were clearly visible, bound to domains 1A, 2A, and 2B of UL5, with the 5′ and 3′ ends bound to domains 2A and 1A, respectively, in agreement with other superfamily I and II helicases^8^. We tentatively modeled them as oligo-dT. Notably, no density of AMP-PNP was observed at the ATPase active site of UL5 (discussed later). The PM consists of the primase domain of UL52 (PD, residues 409-890) and the entire UL8 protein, which interact with each other. The intimate interaction between UL5-UL52 and UL52-UL8 explains why a lack of one subunit seriously affects the helicase and primase activities of the remaining subunits.

**Figure 1:**
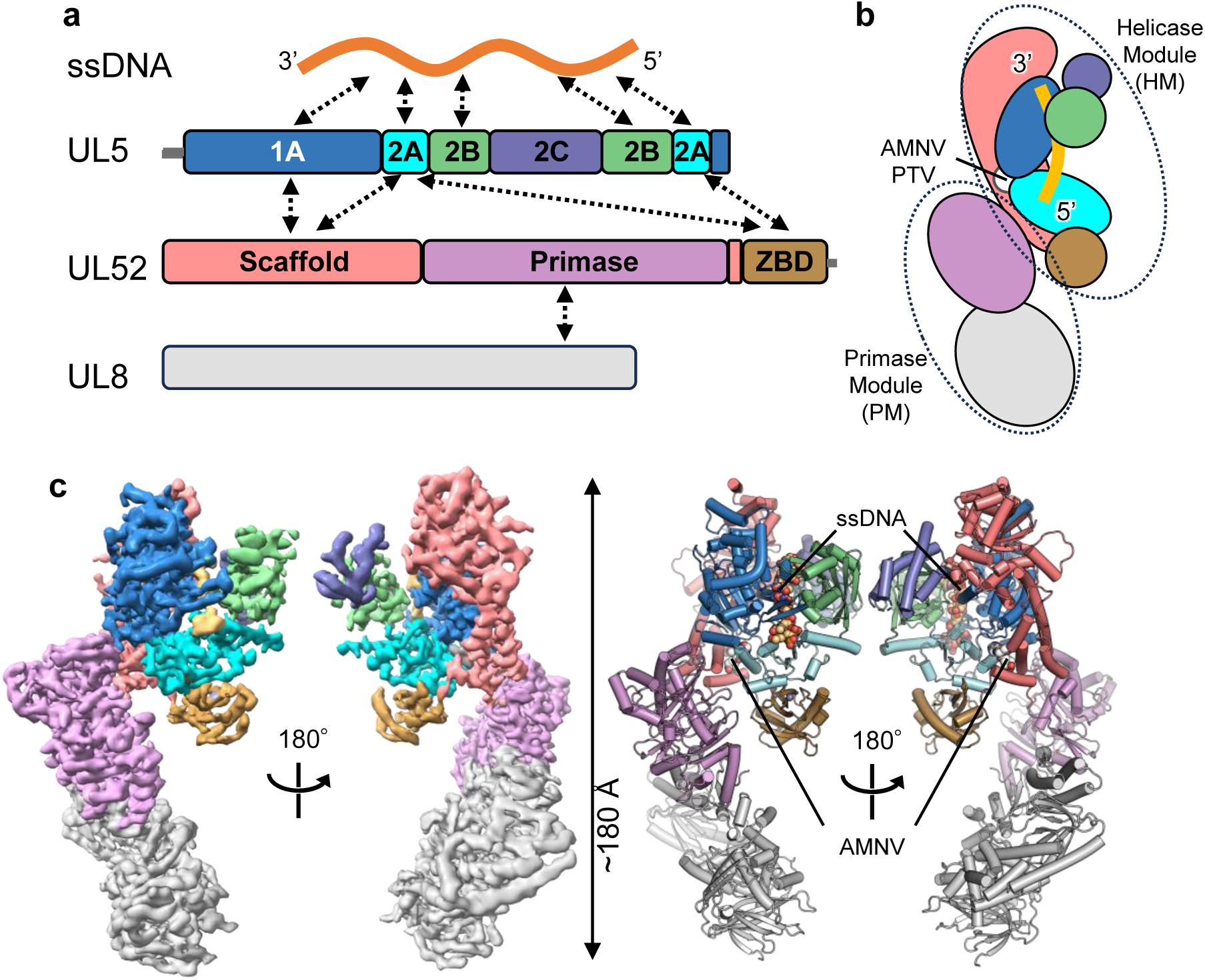
Overall structure of the AMNV-bound HPC from HHV1. a, Domain architectures of the three subunits and their interactions with each other and with ssDNA. The color scheme used in this panel is applied consistently throughout the paper to distinguish different domains. b, A cartoon illustration of the HPC. c, 3DFlex map (calculated using 3DFlex) and the ribbon model of the AMNV-bound HPC. See also Figure S1-S8 and Table S1-S2.

AMNV and PTV bind to a shared pocket located near, but not at, the ATP-binding site. The pocket is formed by the SD of UL52 and the linker region connecting domains 1A and 2A of UL5 (designated as motif IIIa in superfamily I, although sometimes referred to as motif IV)^8^. In the structures of superfamily Ia and Ib helicases^9,20,22^, a conserved arginine residue of motif IIIa (corresponding to R345 of UL5) directly interacts with the γ-phosphate group of the bound ATP analogs. This suggests that motif IIIa plays a role in coupling ATP binding/hydrolysis to domain movement—a process potentially influenced by binding of HPIs. The two inhibitor-bound structures are nearly identical (the RMSD value of 0.50 Å for 1,098 Cα atoms of the HM; Figure S6), suggesting that the two inhibitors likely function in essentially the same manner. Most of the known single-point resistance mutations to AMNV or PTV map directly to this inhibitor-binding pocket, suggesting their physiological binding (see “Binding and specificity of HPIs”).

### PD and ZBD of UL52

The DALI server search revealed that the PD is structurally similar to other archaeo-eukaryotic primases, including human and bacterial PrimPol (e.g., PDB ID: 5L2X; Z-score = 14.8) as well as viral helicase-primase fusion proteins (e.g., PDB ID: 8XJ6; Z-score = 12.7) whose helicase domains assemble into a hexameric ring. When a template DNA-bound structure of PrimPol^23^ is superposed, the superposed DNA is positioned with its 3’ end directed towards the HM, while the HM-bound ssDNA in the current HPC structure is oriented with its 5’ end toward the PM (Figure S7a-c). This configuration is consistent with a model in which a single HPC molecule engages the continuous template strand via its HM and PM simultaneously. However, to accommodate a single ssDNA spanning the two modules in a smooth conformation, substantial reorientation between these two modules would be necessary. Given the apparent flexibility between the two modules and the lack of a large interaction surface between them, it is possible such reorientation occurs during the productive primer synthesis by the PD coupled with replication fork progression driven by the HM. The HPC may engage the template DNA through an alternative mechanism (e.g., two HPC molecules binding to a single replication fork), as suggested earlier^24^.

The ZBD coordinates a zinc ion via C988, H993, C1023, and C1028. The UL52 ZBD is widely conserved among eukaryotic PrimPol proteins. Structural prediction using AlphaFold2^25^ revealed a strong similarity between the ZBD of UL52 and that of human PrimPol (Figure S7d, e). The human PrimPol ZBD possesses ssDNA-binding activity and is required for primase activity. It has been proposed to bind template ssDNA to regulate the primase and polymerase activities of PrimPol^26^. The reorientation of the HM and PM domains may bring the PD and ZBD closer together, enabling such a cooperative binding mode.

### UL8-PD interactions

As predicted by a previous bioinformatic analysis^27^, UL8 adopts a fold that is structurally related to B-family DNA polymerases. However, UL8 lacks both the catalytic residues required for coordinating divalent ions and the basic surface necessary for template DNA binding (Figure S8), consistent with its lack of detectable enzymatic activity. At the UL52–UL8 interface, a network of salt bridges and hydrogen bonds is formed, involving the R640 side chain and L748 main chain of UL8, and the side chains of E511 and R437 of UL52 (Figure S8c). Notably, R640 is one of five residues that are completely conserved among the eight human herpesviruses and was previously predicted to be involved in UL52 binding based on a mutagenesis study^28^.

### Helicase module and ssDNA binding

The bound ssDNA interacts with the domains 1A, 2A, and 2B of UL5. Domains 1A and 2A engage the DNA backbone mainly through hydrophilic interactions with the phosphate groups, while domain 2B mostly forms van der Waals contacts with it (Figure 2a-e). Nucleotides dT3-dT5 are wrapped around by the three domains and completely buried within a channel that is too small to accommodate the double-stranded DNA (Figure 2a), suggesting that like other superfamily I helicases, UL5 unwinds the DNA duplex via a steric exclusion mechanism coupled with the translocation along the tracking strand.

**Figure 2:**
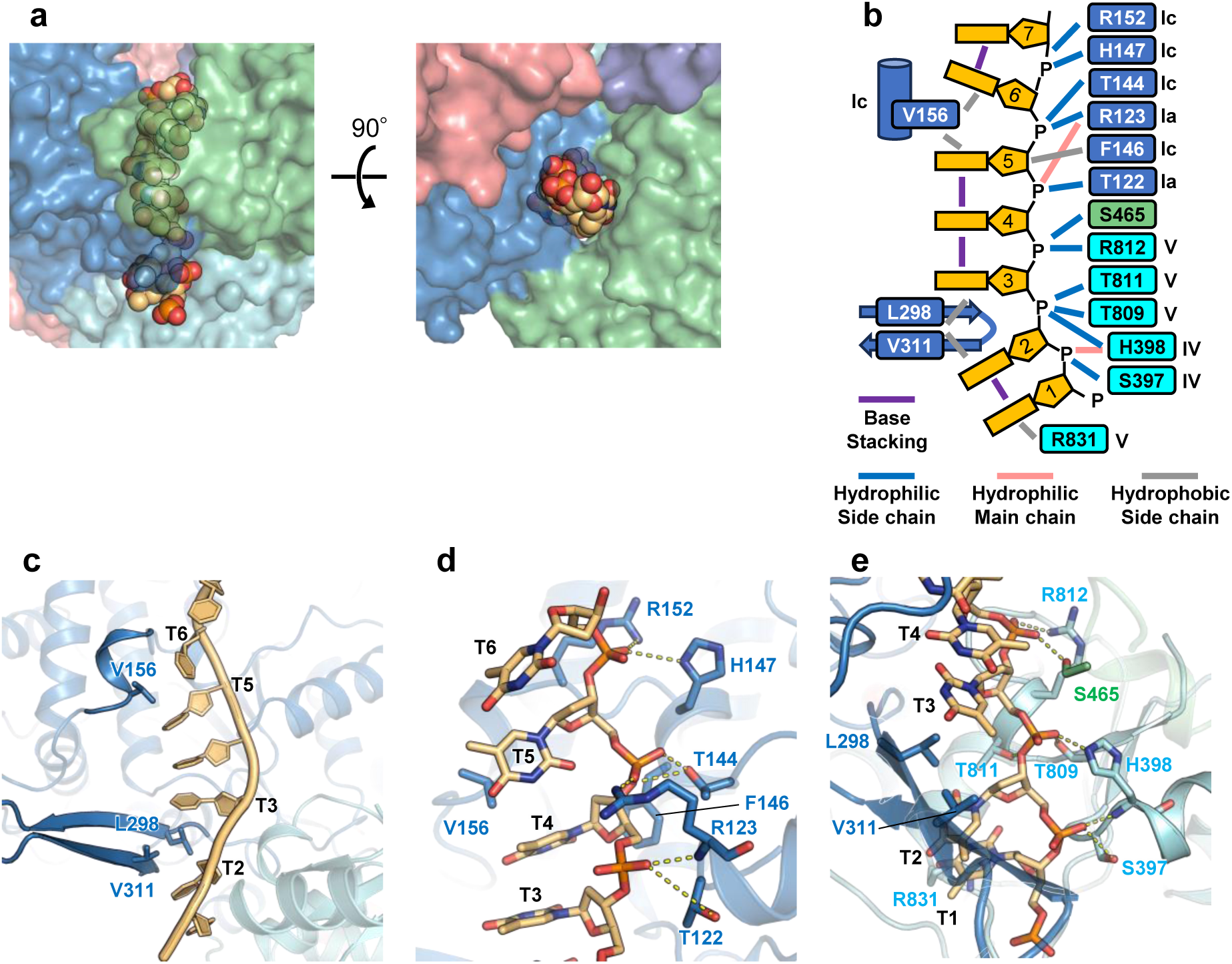
UL5-DNA interaction. a, The bound ssDNA is wrapped by domains 1A, 2A, and 2B (two views). UL5 and ssDNA are shown as a transparent surface model and a sphere model, respectively. b, Schematic representation of ssDNA-UL5 interaction. c, Two structural motifs interfering with base-stacking interactions. d and e, Interactions between ssDNA and domain 1A (d) or domain 2A (e).

Domain 2A binds the phosphate groups of dT2-dT4 via the conserved helicase motifs IV and V, whereas domain 1A binds those of dT5-dT7 via motifs Ia and Ic. This binding mode resembles those observed in other superfamily Ib helicases such as RecD2 and Pif1^9,18–20^. Substitution of T809 (motif V) with isoleucine was shown to impair viral DNA replication and reduce both ATPase and helicase activities of the UL5-UL52 subcomplex^29,30^.

The bound DNA is slightly curved, and base-stacking interactions are interrupted between dT2 and dT3 and between dT5 and dT6 (Figure 2b-e). Two structural elements, the motif Ic helix (residues 153-156) in domain 1A and a β-hairpin (residues 298-313, unique to UL5) of domain 2A, are inserted between the neighboring bases at these positions, respectively, likely stabilizing the curved conformation. Interrupted base stacking is commonly seen in superfamily I and II helicases and may have an important role in DNA unwinding^8^.

The platform domain of UL52 is L-shaped and mostly α-helical. The DALI^17^ search found no hit with comparable side and Z-score > 6.5, indicating that the PD is likely a novel fold. The PD interacts with the UL5, with a large interaction area, and with inhibitors (Figure 1c).

### Domains 1A and 2A adopt an open form in the HPI-bound HPC

During their catalytic cycle, superfamily I helicases undergo conformational changes between an ATP-bound closed form and an ATP-free open form ^8,22^. The ATPase site comprises motifs I (also known as P-loop or Walker A motif), Ia, II (also known as DEXX motif or Walker B motif), III, IIIa, V, and VI, which are distributed along the cleft between domains 1A and 2A. ATP binding induces closure of this cleft, resulting in the formation of a functional ATPase site contributed by both domains. The essential roles of these residues, including the motif I lysine (K103 of UL5), the motif II aspartate and glutamate (D249 and D250 of UL5), and the motif VI arginine (R841 of UL5) have been demonstrated by mutational studies on UL5^29,30^ or related superfamily I helicases^18,31^. Structural studies of superfamily Ia and Ib helicases^9,22^ have shown that motif I and motif II are involved in binding the triphosphate moiety and the catalytic magnesium ion, respectively, whereas the arginine in motif VI functions as the so-called “arginine finger”^8^, contacting the triphosphate moiety in the ATP-bound closed form.

To gain insights into the mode of action of HPIs, we compared the current AMNV-bound HPC structure with that of the structurally most related helicase, Pif1^20^. In both the AMNV-bound HPC and the open form of Pif1, motifs Ia (domain 1A) and IV (domain 1B) contact the bound ssDNA phosphates that are separated by three nucleotides (Figure 3a, b). In contrast, in the closed form of Pif1, the distance between these motifs corresponds to two nucleotides (Figure 3c). This structural comparison clearly shows that the current HPC structures represent an open form. Since the structure of ATP-free HPC in the absence of HPIs has not yet been determined, it remains unclear how closely the current HPI-bound structures resemble the functional ATP-free form in the catalytic cycle.

**Figure 3:**
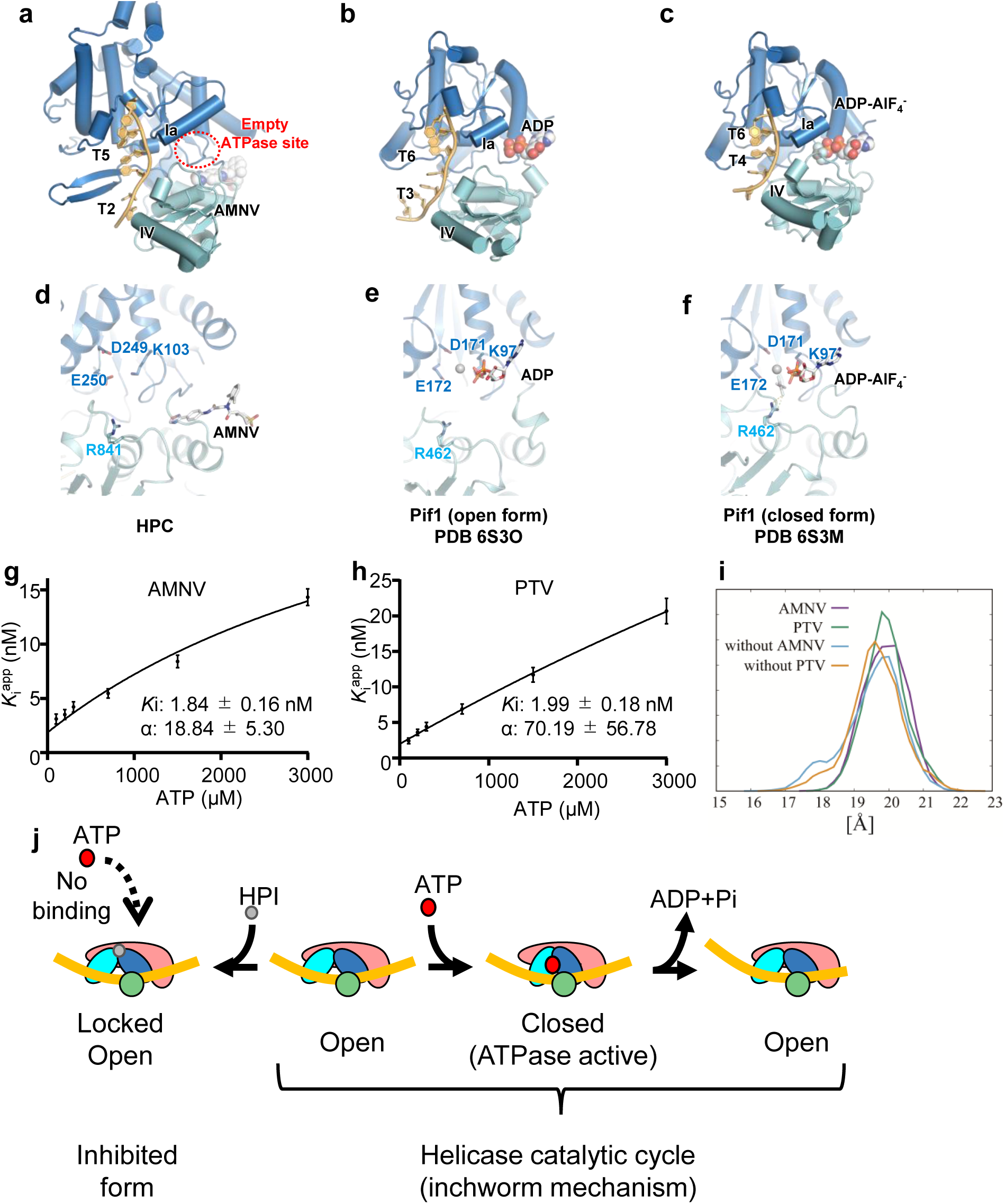
HPIs may lock UL5 in an open conformation. a-c, Structures of domains 1A and 2A in the ssDNA-bound states of the AMNV-bound HPC (a), ADP-bound Pif1 (b), and ATP analog-bound Pif1 (c). DNA nucleotides contacted by conserved residues in motifs Ia and IV (T122 and S397 of UL5 and their equivalents) are labeled. d-f, Close-up views of the ATPase site in the AMNV-bound HPC (d), ADP-bound Pif1 (e), and ATP analog-bound Pif1 (f). g, h, Relationship between ATP concentration and apparent inhibition constant (Kiapp), along with their standard error of the mean (SEM), in ATPase inhibition assays of AMNV (g) and PTV (h). The corresponding Ki and α values with SEM are also shown. i, Distribution of the distance between the centers of mass of K103, D249, and E250 in domain 1A and R841 in domain 2A during MD simulations. j, Proposed model for the inhibition mechanism of HPIs. See also Figure S9 and S10.

We also compared the conformations of the ATPase site among the three structures (Figure 3d-f). In the closed form of Pif1, the bound ATP analog (ADP-AlF₄⁻) is surrounded by residues from both domains 1A and 2A, including residues from motifs I (K97, contacting phosphates), II (D171 and E172, chelating Mg²⁺), and VI (R462, contacting the γ-phosphate analog). In contrast, in the open form of Pif1, the two domains are separated, and the side chain of R462 is too distant to contact the phosphate groups of the ADP bound to domain 1A. Similarly, in the AMNV-bound HPC structures, the corresponding catalytic residues from motifs I (K103), II (D249 and E250), and VI (R841) are also too far apart to coordinate ATP binding. Notably, although 1 mM AMP-PNP was added to the cryo-EM samples, no density was observed at the ATPase site, suggesting that the current open form of HPC has reduced affinity for ATP.

### HPIs allosterically and competitively inhibit ATP binding

The open conformation observed in the HPI-bound HPC structures, along with the absence of AMP-PNP density at the ATPase site, suggests that HPI binding alters the conformation of the HPC, thereby allosterically reducing its affinity for ATP. To test this hypothesis, we performed ATPase inhibition assays (Figure S9) using varying concentrations of ATP and HPIs, and measured the effect of ATP concentration on the apparent inhibition constant (Ki^app^). As shown in Figure 3g and h, increasing ATP concentrations led to a corresponding increase in Ki^app^, suggesting that ATP and HPI binding are reciprocally inhibiting with each other with respect to HPC affinity. We analyzed the inhibition data assuming a mixed inhibition model, in which HPIs can bind both to the free HPC and to the HPC–ATP complex^32^. From this analysis, we calculated the α factor, which represents how differently inhibitors bind to the free HPC versus the HPC–ATP complex. Curve fitting yielded large α factor values for both inhibitors (about 19 for AMNV and 70 for PTV), indicating that both inhibitors bind to the free enzyme much more strongly, acting in a nearly ATP-competitive manner, despite binding at the allosteric site.

### HPIs may lock the HPC in the open form

Our structural and biochemical analyses suggest that HPIs affect the conformational dynamics of the HPC by locking it in the open conformation observed in our cryo-EM structures. To further substantiate our findings, we performed molecular dynamics (MD) simulations on the following four systems (Figure S10a). The first two systems consist of the UL5-UL52 subcomplex bound to ssDNA, with either AMNV or PTV (based on the AMNV- or PTV-bound HPC structures, respectively). The remaining two systems are identical to the first two, but lack the inhibitors, serving as controls.

For each system, we conducted eight replica MD simulations, each lasting 3 µs, and analyzed the distance between key catalytic residues in domains 1A and 2A: K103, D249, and E250 from motifs I and II in domain 1A, and R841 from motif VI in domain 1B (Figure 3d). As shown in Figure 3i, the HPI-bound systems exhibited a unimodal distribution of inter-residue distances, peaking around 20 Å. In contrast, the HPI-free systems showed a bimodal distribution with peaks centered at approximately 18 Å and 19.5 Å. These results suggest that, in the absence of HPIs, domains 1A and 2A undergo spontaneous conformational fluctuations that may potentially facilitate productive ATP binding and hydrolysis. In contrast, HPI binding appears to restrict this flexibility, stabilizing the HPC in an open, catalytically inactive conformation.

Based on these findings, we propose the following inhibition model (Figure 3j). In the catalytic cycle of the inchworm model, the helicase alternates between open and closed conformations, driven by ATP binding and hydrolysis. However, upon HPI binding, the HPC is locked in the open conformation, thereby preventing ATP binding and stalling the catalytic cycle. Consequently, the helicase remains inactive and unable to unwind DNA.

### Binding and specificity of HPIs

Figure 4a-c shows the interactions between AMNV and PTV with the HPC, respectively. In both complexes, two connected aromatic rings of each HPI are surrounded by residues N98, N343, G352, M355, and K356 of UL5, as well as Y356 of UL52. AMNV and PTV contain two and one carbonyl group(s), respectively, in their central regions, which are within hydrogen-bonding distance of the UL5 K356 side chain. A sulfonyl group, which is shared by both HPIs, forms a hydrogen bond with the side chain of UL52 N902. The dimethylphenyl group, unique to AMNV, engages in hydrophobic interactions with the side chains of UL5 Y882 and UL52 F360. Notably, most of the known single-point resistance mutations of AMNV (G352C, M355T, and K356N in UL5, as well as F360V/C and N902T in UL52)^33^ and PTV (N342K, G352V, M355T, and K356Q in UL5 as well as A899T in UL52)^14,34,35^ are located at or near the binding site. N342K, G352C/V, and M355T in UL5 are expected to cause steric clashes at the HPI binding site, which may hinder HPI binding. K356N/Q in UL5, as well as F360V/C and N902T in UL52, are predicted to disrupt critical HPC–HPI interactions. These structural alterations likely account for their resistance mechanisms. Although the HPI-contacting residues in UL5 are well conserved among all human herpesviruses, those in UL52, such as F360, A899, N902, H906, and F907, are less conserved (Figure 4d). This observation explains why the current HPIs are ineffective against β- and γ-herpesviruses and provides a structural basis for the rational design of novel HPIs targeting these subfamilies.

**Figure 4:**
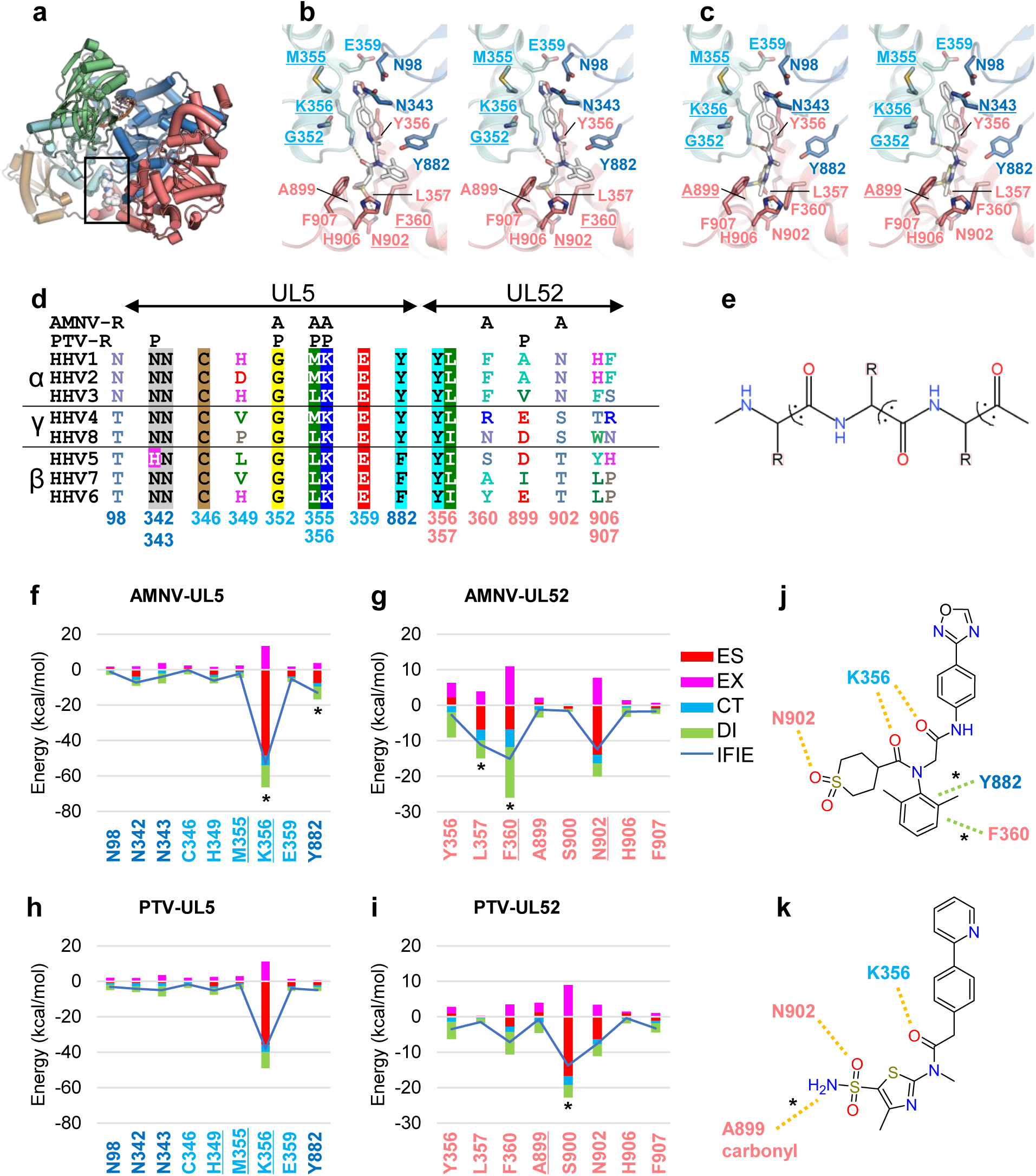
Binding and specificity of HPIs. a, Overall structure of the HPC, with the HPI binding site highlighted by a box. b, c, Stereo views of the binding site for AMNV (b) and PTV (c). Residues associated with resistance mutations are underlined. d, Sequence alignment of residues involved in HPI binding. “A” and “P” denote mutations that confer resistance to AMNV and PTV, respectively. e, Fragmentation scheme used for FMO calculations. f–i, PIEDA results showing interactions between AMNV and UL5 (f), AMNV and UL52 (g), PTV and UL5 (h), and PTV and UL52 (i). ES, EX, CT, and DI refer to the electrostatic, exchange-repulsion, charge transfer with higher-order mixing, and dispersion terms, respectively. Asterisks indicate residues contributing more than 5 kcal/mol to binding in one complex compared to the other. j, k, Schematic representations of the interactions between the HPC and AMNV (j) or PTV (k). See also Figure S11.

We next analyzed the MD trajectories to assess the potential contribution of water molecules to the HPI binding. As shown in Figure S10b and c, a region of high water density was observed in both complexes, where two water molecules bridge the side chain of UL5 K356 and the sulfonyl group of the HPIs. Notably, these bridging water molecules connect UL5 K356 and the sulfonyl group for 39.1% and 45.0% of the simulation time in the AMNV and PTV complexes, respectively, suggesting the formation of a stable hydrogen-bonding network. This water-mediated interaction may explain why the sulfonyl group is essential for the tight binding of these inhibitors.

To quantitatively evaluate the differences between the AMNV and PTV-bound structures and to gain mechanistic insights into their distinct anti-herpesvirus spectra, we performed fragment molecular orbital (FMO) calculation analysis^36–39^ on the representative structures of MD trajectories. The FMO method divides biological macromolecules into residue-based fragments (Figure 4e), thereby accelerating quantum chemical calculations and enabling the extraction of inter-fragment interaction energy (IFIE) and its decomposition data. To account for structural fluctuations in the HPI-HPC interactions, we clustered the MD trajectories into four groups based on the coordinates of the HPI binding site (Figure S10d), performed FMO calculations on the representative structures, and calculated the weighted average energies, taking into account the existence probabilities of each cluster.

Figure 4f-i summarizes the FMO analysis results, showing the frequency-weighted average values of IFIE and pair interaction energy decomposition analysis (PIEDA)^40^ between either AMNV or PTV and the UL5 and UL52 subunits. In agreement with our manual inspection, residues involved in recognizing the common structural features of the two HPIs are consistently identified in both complexes. These include residues surrounding the two connected aromatic rings (UL5 N98, N343, and M355), UL5 K356 forming hydrogen bonds with the HPIs’ carbonyl oxygen, and UL52 N902 forming a hydrogen bond with the sulfonyl oxygen.

In addition, comparison of the IFIE and PIEDA data (Figure 4f-i) reveals distinct binding features of AMNV and PTV (summarized in Figure 4j, k). The stronger attractive interaction energies for UL5 Y882 and UL52 F360 in the AMNV complex (by approximately 8.1 and 7.9 kcal/mol, respectively), consisting of considerable dispersion interaction energy (Figure 4f-i), indicate that the recognition of AMNV’s unique dimethylphenyl group by these residues via π-π and CH/π interactions, respectively (Figure 4b), substantially contributes to the binding affinity. Notably, these residues are conserved across the α-subfamily, consistent with the AMNV’s broad efficacy against all α-subfamily members (Figure 4d). In contrast, UL52 S900 exhibits a stronger IFIE value in the PTV complex, mainly due to electrostatic and charge transfer interactions, suggesting the presence of the hydrogen bond^39^. In the FMO method, proteins are fragmented between the α-carbon and carbonyl carbon, assigning the carbonyl group to the next residue (+1) (Figure 4e). Structural analysis of the PTV complex revealed a hydrogen bond between the A899 carbonyl group (assigned as S900 in the FMO scheme) and the PTV-specific NH₂ group of the sulfonamide moiety (Figure S11a). Structural modeling suggests that the substitution of A899 with threonine (which confers PTV resistance) or with valine (the corresponding residue in HHV3, Figure 4d) introduces steric hinderance that disrupts this interaction (Figure S11b, c). Moreover, the A899T substitution does not confer resistance to BILS 22 BS, another HPI that retains a sulfonyl group but lacks the NH₂ group^35^.

Collectively, these findings indicate that the unique hydrogen bond between the NH₂ group of PTV and the A899 carbonyl oxygen is critical for its inhibitory activity, and explain PTV’s limited activity against HHV3.

## Discussion

Our structural, biochemical, and computational analyses collectively elucidate the mechanism of action of AMNV and PTV and shed light on why these HPIs are less effective against the β- and γ-herpesvirus subfamilies. Moreover, our MD and FMO analyses clarify how a small difference in the chemical structures of AMNV and PTV results in distinct anti-herpesvirus spectra within the α-subfamily. Together, these findings establish a structural framework for the rational design of novel HPIs with tailored anti-herpesvirus spectrum and/or improved pharmacokinetic properties.

Structures solved by cryo-EM often lack information on conformational dynamics, and may suffer from limited resolution, leading to uncertainty in positioning of ligands and surrounding water molecules. To overcome these limitations, we performed molecular dynamics (MD) simulations to capture structural fluctuations and water behavior, followed by FMO analysis to quantitatively evaluate molecular interactions. This integrated approach provided in-depth insights into HPI–HPC interactions that aligned well with previous mutational studies, supporting the reliability of our methodology. The combined use of MD and FMO thus represents a powerful framework for investigating systems where structural data alone is insufficient to fully characterize essential molecular interactions^41,42^.

Helicases, which play essential roles in nearly all aspects of nucleic acid metabolism, have long been recognized as promising drug targets. However, developing effective helicase inhibitors has proven challenging, likely due to the chemical nature of the ATP- and nucleic acid-binding sites, which are difficult to target with small molecules^43,44^. Recent studies have reported the development of specific helicase inhibitors that act allosterically^45–47^. In this study, we demonstrate that AMNV and PTV, two clinical inhibitors, bind to a shared allosteric pocket at the interface between the helicase subunit (UL5) and the non-helicase subunit (UL52). Our detailed characterization of these clinical HPIs, which act allosterically yet competitively, provides valuable insights for the development of diverse inhibitors targeting helicases from a wide range of sources, including both cellular and pathogenic origins.

The current HPC structures shed light on the coupling mechanism between DNA unwinding and primer synthesis; however, several questions remain unresolved. Do UL5 and UL52 engage with a single continuous ssDNA strand? How do the HM and PM reposition themselves during the catalytic cycle? Does the ZBD participate in binding the ssDNA template to facilitate efficient primer synthesis by the PD? To address these questions and elucidate the coupling mechanism, it will be necessary to determine the structures of HPC actively engaged in both DNA unwinding and primer synthesis in both ATP-bound and ATP-free states. Such structural insights will also be valuable for deepening our understanding of DNA replication mechanisms across diverse organisms.

## Resource availability

### Lead contact

Further information and requests for resources and reagents should be directed to and will be fulfilled by the lead contact, Toru Sengoku.

### Materials availability

Plasmids generated in this study are available from the lead contact upon request.

### Data and code availability

Coordinates and density maps have been deposited in Protein Data Bank (accession codes 9UT1, 9UT3, 9UT4, 9UT5, 9UT6, and 9UT7) and Electron Microscopy Data Bank (accession codes EMD-64478, EMD-64479, EMD-64480, EMD-64481, EMD-64482, and EMD-64483), respectively. All structure files and a set of input/output files used for FMO calculations are available at the FMODB (https://drugdesign.riken.jp/FMODB/)^48^ The FMODBIDs for the representative structures of AMNV-HP complex in clusters 0-3 are YJ552, 5817Z, 4812N, and KM4Y3, respectively. The FMODBIDs for the representative structures of PTV-HP complex in clusters 0-3 are QJ2RY, RV7Z8, Z14JN, and 681MZ, respectively. Simple data analysis can be performed using the FMODB web interface, and detailed analysis can be performed using the BioStation Viewer software (https://fmodd.jp/biostationviewer-dl/).

## Supporting information

Overall structure of the AMNV-bound complex

AMNV binding site

## Acknowledgements

We thank members of Nureki Laboratory for their support in cryo-EM data collection. MD calculations were performed using computational resources of the supercomputer system (HPE SGI8600) jointly operated by the Japan Atomic Energy Agency (JAEA) and the National Institutes for Quantum Science and Technology (QST). FMO calculations were performed as part of the activities of the FMO Drug Design Consortium (FMODD) using the supercomputer “Fugaku” at RIKEN (project ID: hp240162).

This work was supported by JSPS KAKENHI (grant number 25K02501), Sumitomo Foundation, Shionogi Infectious Disease Research Promotion Foundation, and Uehara Memorial Foundation to T.S. This research was also partially supported by Research Support Project for Life Science and Drug Discovery (Basis for Supporting Innovative Drug Discovery and Life Science Research (BINDS)) from Japan Agency for Medical Research and Development (AMED) under grant numbers JP25ama121012 (support No. 6021) to O.N., JP25ama121024 (support No. 5660) to H.K., and JP25ama121030 (Support No. 5726) to K.F.

## Author contributions

K.S. purified proteins, collected the cryo-EM data, solved the structures, performed biochemical experiments, analyzed data, and wrote the paper. H.I. and H.K. performed MD simulations and wrote the paper. T.M. and K.F. performed FMO analysis and wrote the paper. S.K. purified proteins and performed biochemical experiments. Y.K., K.H., and O.N. assisted with cryo-EM data collection. C.O. and A.O. purified proteins. T.S. designed the project, solved the structures, analyzed data, and wrote the paper.

## Declaration of interests

The authors declare no competing interests.

## Declaration of Generative AI and AI-assisted technologies in the writing process

During the preparation of this work the author used ChatGPT in order to improve readability and language. After using this service, the authors reviewed and edited the content as needed and take full responsibility for the content of the publication.

## Methods

### Experimental model and study participant details Cell lines

Spodoptera frugiperda (Sf9) insect cells were cultured and maintained in Sf-900 II SFM insect cell culture medium (Thermo Fisher Scientific) at 27 °C.

### Methods details

#### Protein expression and purification

Full-length coding sequences of HHV-1 UL5, UL52, and UL8 were synthesized by Twist Bioscience and individually cloned into the pACEBac1 vector. An N-terminal twin-strep–His tag was fused to UL5, while N-terminal His–SUMOstar tags were fused to UL52 and UL8. Baculoviruses were generated in Sf9 cells using the Bac-to-Bac system (Thermo Fisher Scientific) according to the manufacturer’s protocol. The HPC was reconstituted by co-infecting Sf9 cells with the three respective recombinant baculoviruses at a multiplicity of infection of 3:3:3. Cells were harvested 48 hours post-infection.

Cell pellets were resuspended in lysis buffer (100 mM Tris-HCl (pH 8.0), 150 mM NaCl, 1 mM MgCl₂, 0.5 mM TCEP, 0.5% Tween 20, 10% glycerol, 0.1 mM PMSF, and cOmplete ULTRA protease inhibitor cocktail (Roche)) and lysed by sonication. The clarified lysate was loaded onto a Strep-Tactin XT 4Flow column (IBA Lifesciences), and the bound protein complex was eluted with buffer containing 20 mM Tris-HCl (pH 8.0), 500 mM NaCl, 10% glycerol, 0.5 mM TCEP, and 50 mM D-biotin. The eluate was subsequently applied to a HisTrap HP column (Cytiva) and eluted with buffer containing 20 mM Tris-HCl (pH 8.0), 500 mM NaCl, 10% glycerol, 0.5 mM TCEP, and 500 mM imidazole. The eluted sample was dialyzed against buffer A (20 mM Tris-HCl (pH 8.0), 150 mM NaCl, 0.5 mM TCEP) and concentrated for cryo-EM sample preparation.

For tag removal, the HisTrap eluate was mixed with SUMOstar protease and HRV 3C protease and dialyzed against buffer A. The sample was subsequently passed through a HisTrap column to remove cleaved tags and proteases. The flow-through containing tag-free HPC was further dialyzed against buffer A, concentrated, flash-frozen in liquid nitrogen, and stored at −80 °C for ATPase assays.

### Cryo-EM sample preparation

For grid preparation of the AMNV-bound complex, 0.4 μM tagged HPC was mixed with 4 μM 43-mer ssDNA (5’-TTTTTTTTTCCTGTTTTTTTTTTTTTTTTTTTTTTTTTTTTTT- 3’; Fasmac), 1 mM AMP-PNP, 10 mM MgCl2, and 100 μM AMNV (AstaTech), and then concentrated 10-fold. For the PTV-bound complex, 0.48 μM tagged HPC was mixed with 4.8 μM 43-mer ssDNA, 1 mM AMP-PNP, 10 mM MgCl2, and 100 μM PTV (Selleck), and then concentrated 7.6-fold. Concentration was performed using a Vivaspin 500 device (MWCO 50 kDa; Sartorius).

Due to orientation bias observed on Au 300-mesh R1.2/1.3 grids, the samples were applied to freshly glow-discharged Au 300-mesh R0.6/1 holey carbon grids (Quantifoil), pre-treated with 3 μL of amylamine to mitigate bias^49^. The grids were blotted for 4 s at 4 °C under 100% humidity and plunge-frozen in liquid ethane using a Vitrobot Mark IV (Thermo Fisher Scientific).

### Cryo-EM data acquisition and processing

Cryo-EM data were collected using a Titan Krios G4 transmission electron microscope (Thermo Fisher Scientific) operated at 300 kV, equipped with a K3 direct electron detector (Gatan), installed at the University of Tokyo. Data acquisition was performed using EPU v3.4.0.5704, with a defocus range of −0.6 to −1.6 μm. Data acquisition statistics are summarized in Supplementary Table 1.

The cryo-EM data were processed using CryoSPARC v4.4.1^50^ (Figures S1 and S2). For the AMNV complex, an initial dataset of 803 micrographs was used for blob picking, 2D classification, and template generation. These templates were then applied to the main dataset of 12,466 micrographs for template-based particle picking, followed by neural network-based picking using Topaz^51^. Subsequent reconstruction and refinement yielded a global map with poorly resolved density in the HM. Following 3D classification without a focus mask, seven out of ten classes were selected, which showed slightly improved HM density. Particles corresponding to the HM and PM were then subtracted and used for local refinement. The global map was further refined using 3DFlex^16^.

For the PTV complex, data were processed similarly to the AMNV dataset. However, the AMNV-derived templates were reused for template-based particle picking, and no particle subtraction was performed.

### Model building and refinement

The initial model of the HPC was generated using LocalColabFold^52^, which employs AlphaFold2^53^ for structure prediction, and was fitted into the cryo-EM map. The predicted model showed good overall agreement with the map, although notable differences were observed in local loop regions (including the HPIs binding pocket) and in the relative orientation between the HM and PM. The model was manually adjusted using Coot v0.9.8.93^54^ and further refined with the *phenix.real_space_refine* module in Phenix v1.21.1-5286^55^. No map sharpening was applied. To build the atomic models of the HPIs, multiple conformations of their functional groups (the oxadiazole ring of AMNV, and the pyridine ring and NH₂ group of PTV) were generated and evaluated using FMO calculations. The conformation with the lowest energy was then selected. Structural figures were prepared using PyMOL v2.5.0 (Schrödinger, LLC) and ChimeraX v1.9^56^. Data processing and refinement statistics are summarized in Supplementary Table 2.

### ATPase assay

ATPase activity was assessed using an NADH-coupled microplate assay as previously described^57^, performed over a 30-minute time course at 30 °C, with absorbance measurements taken every 30 seconds. Each 54 µL reaction was prepared in ATPase buffer containing 20 mM HEPES-NaOH (pH 7.5), 5 mM MgCl2, 10% glycerol, 5 mM TCEP, 100 µg/mL bovine serum albumin, 2 µM 43-mer ssDNA, 3 mM phosphoenolpyruvate (Nacalai Tesque), 1 mM NADH (Sigma-Aldrich), 1% DMSO, 0.02 U/µL L-lactate dehydrogenase (Sigma-Aldrich), 0.02 U/µL pyruvate kinase (Sigma-Aldrich), 10 nM tag-free HPC, and ATP (Nacalai Tesque) at final concentrations of 0, 0.1, 0.2, 0.3, 0.7, 1.5, or 3 mM. Either AMNV or PTV was added separately at concentrations of 0, 2.5, 5, 10, 15, 20, or 25 nM. Reactions were dispensed into a 96-well half-area microplate (Greiner Bio-One), and NADH oxidation was monitored by measuring absorbance at 340 nm using a Tecan Spark Cyto 400 multimode microplate reader. Reaction velocities (v, OD/sec) were determined from the linear portion of the absorbance decrease at 340 nm between 1000 and 1500 seconds. Background signals measured in the absence of ATP were subtracted to correct the data. The Michaelis constant (*K*M) was determined by non-linear fitting to the Michaelis–Menten equation using GraphPad Prism v10.2.3 (GraphPad Software).

Both AMNV and PTV exhibited EC50 values of less than 100 nM against the virus ^12,14^, indicating their behavior as tight-binding inhibitors ^32,58^. The apparent inhibition constant (*K*ᵢ^ₐₚₚ^) was determined by non-linear fitting to the Morrison equation (Eq. 1), where vi is the initial rate of the inhibited reaction, v0 is the initial rate of the uninhibited reaction, [E] is the enzyme concentration, [I] is the inhibitor concentration, and *K*ᵢ^ₐₚₚ^ is the apparent inhibition constant ^32,58^.

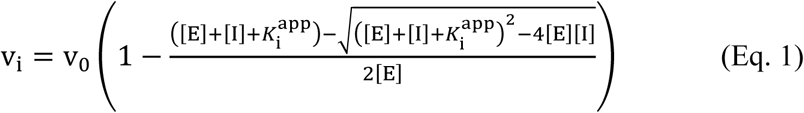

The *K*ᵢ^app^ values were plotted against ATP concentrations and fitted by non-linear regression to a mixed inhibition model (Eq. 2), in which [S] represents the ATP concentration, *K*M is the Michaelis constant, *α* reflects the difference in inhibitor affinity for the free enzyme versus the enzyme-substrate complex, and *K*i is the inhibition constant ^32,58^. This analysis yielded values for α and *K*i.

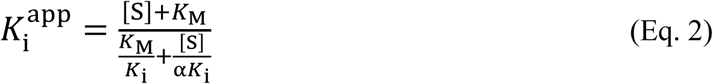

### Modeling of the UL5**–**UL52 subcomplex for MD simulations

For UL5, the N-terminal region (M1–A33) and a large C-terminal loop (A582–E657) were excluded from the model. A shorter missing loop (A563–E568) was modeled using MODELLER^59^, and the termini adjacent to the large loop (T581, T658) were capped with N-methylamide (NME) and acetyl (ACE) groups, respectively. For UL52, the N- and C-terminal regions (M1–T16, P1044–P1058), as well as a large internal loop (P682–P752), were also excluded. The termini flanking this large loop (R681, V753) were capped in the same manner. Three smaller loops (S170–D183, P481–C498, R995–A996) were modeled with MODELLER by generating 100 loop conformations with the remaining structure held fixed; the lowest-energy models were selected for each. The resulting structure was used for subsequent molecular dynamics simulations.

### MD simulations of the UL5**–**UL52 subcomplexes

We prepared four systems of the UL5–UL52 complex: two containing ligands (AMNV or PTV) and two ligand-free systems generated by removing the respective ligands. MD simulations were performed using the pmemd module in AMBER20^60,61^, with force fields ff99SB for protein^62^, bsc1 for DNA^63^, and ff99ions08 for ions^64^. The force field parameters for ligands were prepared using the Antechamber package^65^ in AmberTools and the general Amber force field (GAFF)^66^ for organic molecules. A quantum chemical calculation to geometrically optimize the ligands were carried out with HF/6-31G* basis set using Gaussian 03 software^67^. The net charge for each ligand was set at 0. Discrete partial atomic charges were assigned with HF/6-31G* basis set using the Restrained Electrostatic Potential (RESP) fitting method^68^. The zinc AMBER force field (ZAFF) for the 4-coordinated zinc metal centers^69^ was used for the zinc ion in the ZBD. The UL5 – UL52 complex was solvated in a cubic periodic box of ∼ 190 Å × 190 Å × 190 Å, containing ∼ 220,000 TIP3P water molecules^70^ and 0.15 M NaCl to mimic physiological ionic strength. The complex was positioned such that all atoms were separated at least 20 Å from the edge of the box. The resulting system contained ∼ 690,000 atoms in total.

For each of the four systems, the MD simulations were carried out using eight replicas of identical structures, each initiated with different random seeds for velocity assignment. Each replica was run for 3.16 μs, totaling 25.28 μs per system, under constant pressure of one bar and temperature of 300 K. The dielectric constant was set to 1.0 and the van der Waals interactions were calculated with a 9 Å cutoff. Electrostatic interactions were treated using the particle mesh Ewald (PME) method^71^, with a charge grid spacing close to 1 Å. The charge grid was interpolated using a fourth-order cubic B-spline with the direct sum tolerance of 10^-5^ at a 9 Å direct space cutoff. Temperature and pressure were regulated by Langevin dynamics with a collision frequency of 2 ps^-1^; coupling times were set to 1 ps^-1^ for both. Bond lengths involving hydrogen atoms were constrained with the SHAKE algorithm^72,73^. The leap-frog algorithm was used to integrate the equations of motion with a 2 fs time step. The system was gradually heated from 0 K to 300 K over 10 ns, during which the solute atoms were restrained with decreasing force constants while water molecules and ions were allowed to move freely. Conformational snapshots were saved every 100 ps. Trajectories from the final 3 μs of each of the eight replicas (totaling 24 μs) were used for analysis. Water density maps were generated using the volmap tool in VMD^74^.

### K-mean clustering of the MD trajectories

K-means clustering was performed on all atoms in residues located within 3.5 Å of the ligands in the energy-minimized structure. Specifically, these residues were Asn98, Ile341, Asn343, Met355, Lys356, Glu359, Tyr836, and Tyr882 of UL5, and Tyr356, Phe360, Ala899, and Asn902 of UL52. K-means clustering with K = 4 grouped the trajectories for the UL5-UL52 complex with AMNV into four distinct clusters: 10.5%, 40.2%, 42.7%, and 6.6% of the total trajectory. For the PTV-bound complex, the corresponding cluster populations were 70.0%, 4.9%, 0.1%, and 25.0%. Representative structures were selected as those closest to the centroids of each cluster. The RMSD values relative to the centroids were 0.659, 0.515, 0.588, and 0.703 Å for the AMNV-bound UL5–UL52 complex, and 0.568, 0.555, 0.779, and 0.520 Å for the PTV-bound complex.

### FMO calculation and interaction energy analysis

FMO calculations were performed on the representative structures obtained by clustering of MD trajectories. As a preprocessing step, MD representative structures were subjected to energy minimization by applying a positional constraint of 1.0 kcal/mol-Å^2^ to heavy atoms, and water within 10 Å around the HPC protein, HPI, ssDNA, and ligand was extracted for calculation. Fragmentation was performed by using amino acid residue units for proteins, base and sugar-phosphate fragments for DNA, and single fragments for ligand and water molecules^75^. The FMO calculations were performed at the MP2/6-31G*^76,77^ theoretical level using the ABINIT-MP program^78^. The supercomputer Fugaku was used.

Defining the inter-fragment interaction energy (IFIE) between fragments i and j as 𝐸_𝑖𝑗_, pair interaction energy decomposition analysis (PIEDA)^40^ divides IFIE into four energy components as follows.

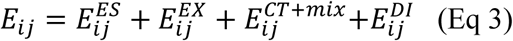

In this study, *i* is HPI (AMNV or PTV), and *j* is each residue of HPC. Weighted average IFIE/PIEDA, 𝐸_ij_^𝑤𝑒𝑖𝑔ℎ𝑡^, is the sum of the interaction energies between HPI and each residue in the cluster representative structure, weighted by the occurrence frequency of each cluster.

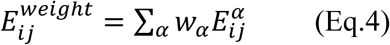

Here, 𝑤_𝛼_ is the weight of cluster α, which is 0.105, 0.402, 0.427, and 0.066 for AMNV, and 0.700, 0.049, 0.001, and 0.250 for PTV. Figure 4f-i shows the weighted average interaction energy calculated using Eq. 4.

Cryo-EM structures were also evaluated using MP2/6-31G* level FMO calculations. For preprocessing of the structures, hydrogen atoms were added and energy minimization was performed using Amber10:EHT force field.

## QUANTIFICATION AND STATISTICAL ANALYSIS

Cryo-EM data collection and refinement statistics are summarized in Table S1.

## Supplemental Information

Document S1. Figures S1-S11, Tables S1, and S2.

Video S1. Overall structure of the AMNV-bound complex.

Video S2. AMNV binding site.

---

**Figure S1:**
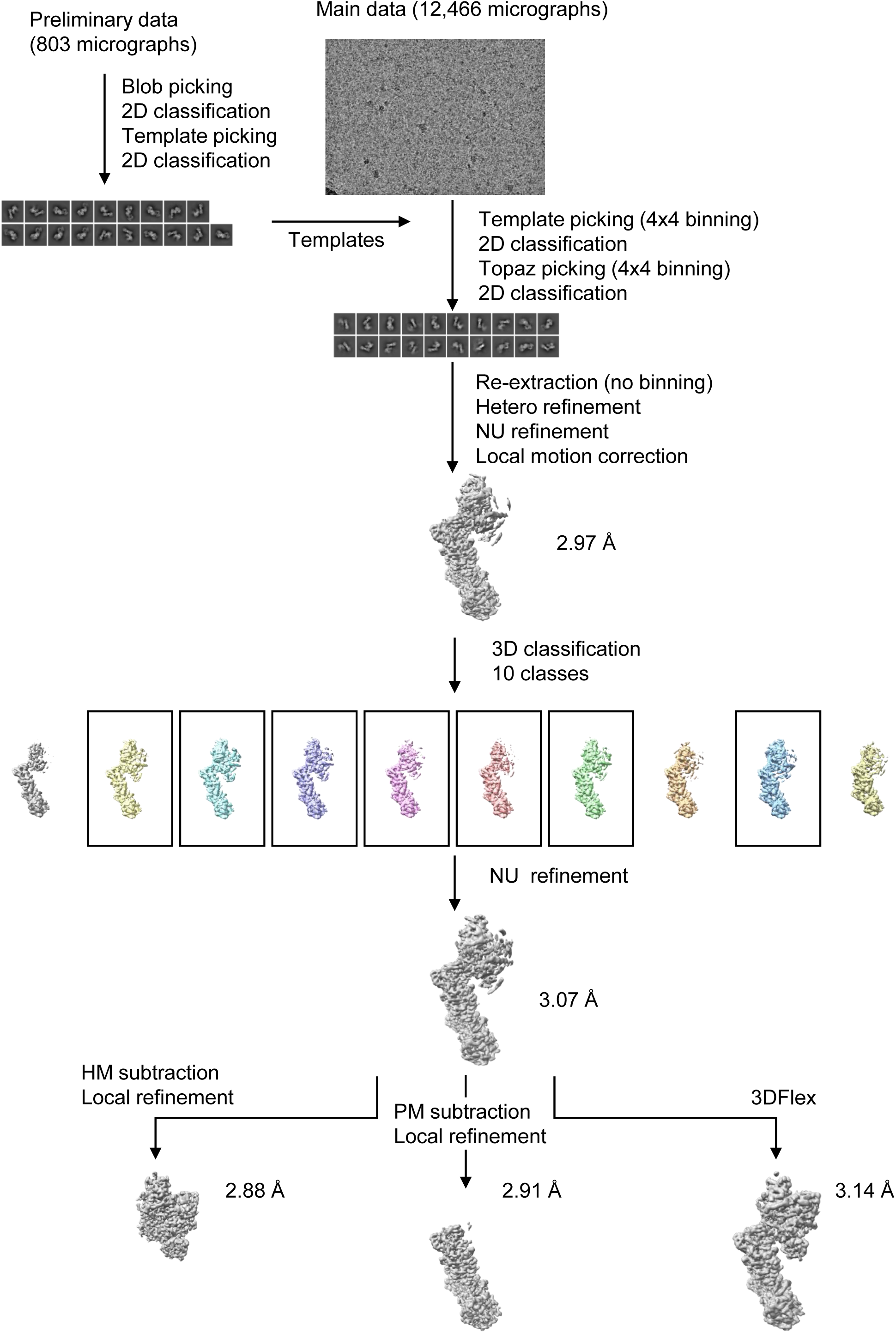
Cryo-EM data processing of the AMNV complex, related to Figure 1.

**Figure S2:**
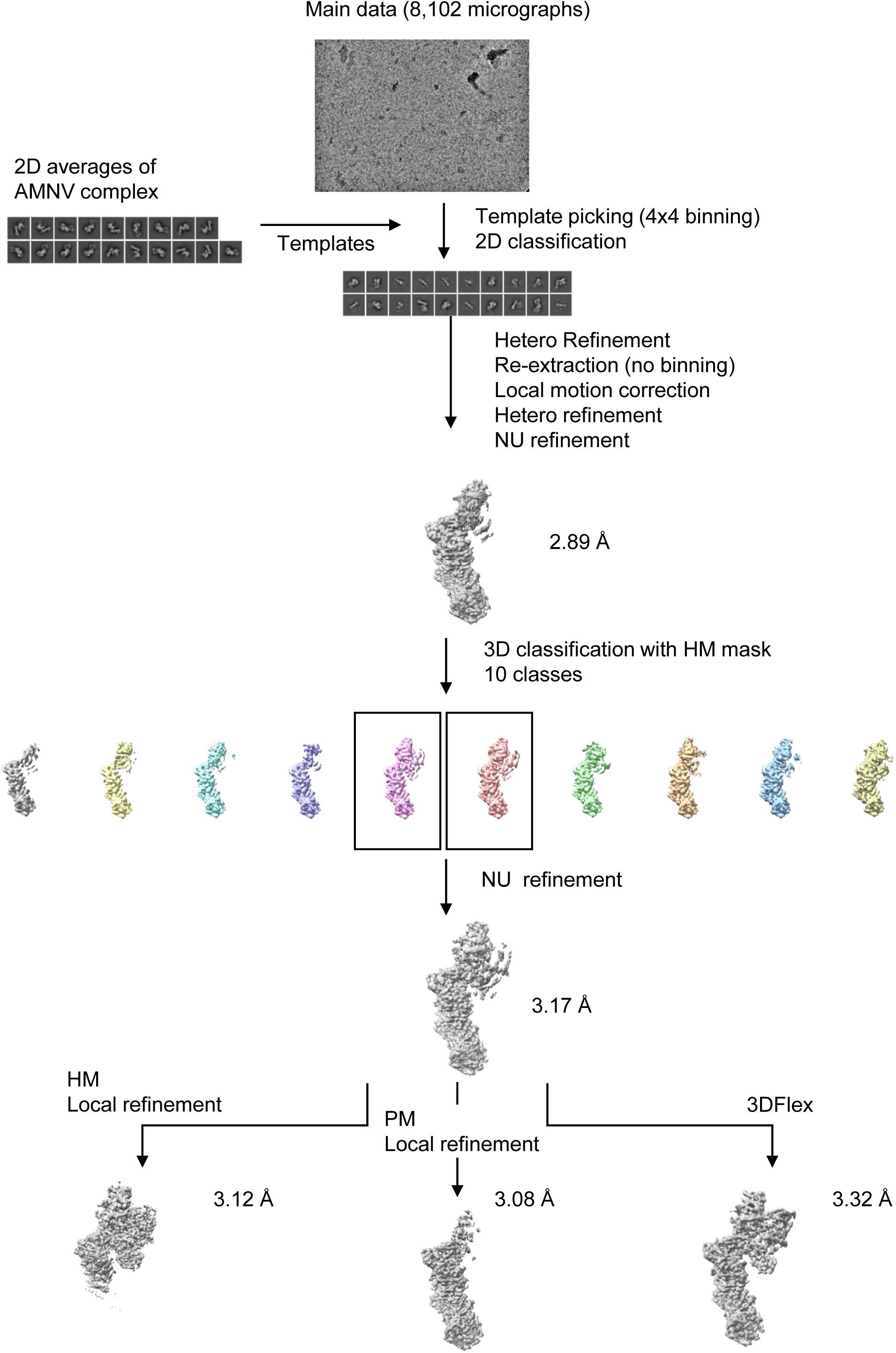
Cryo-EM data processing of the PTV complex, related to Figure 1.

**Figure S3:**
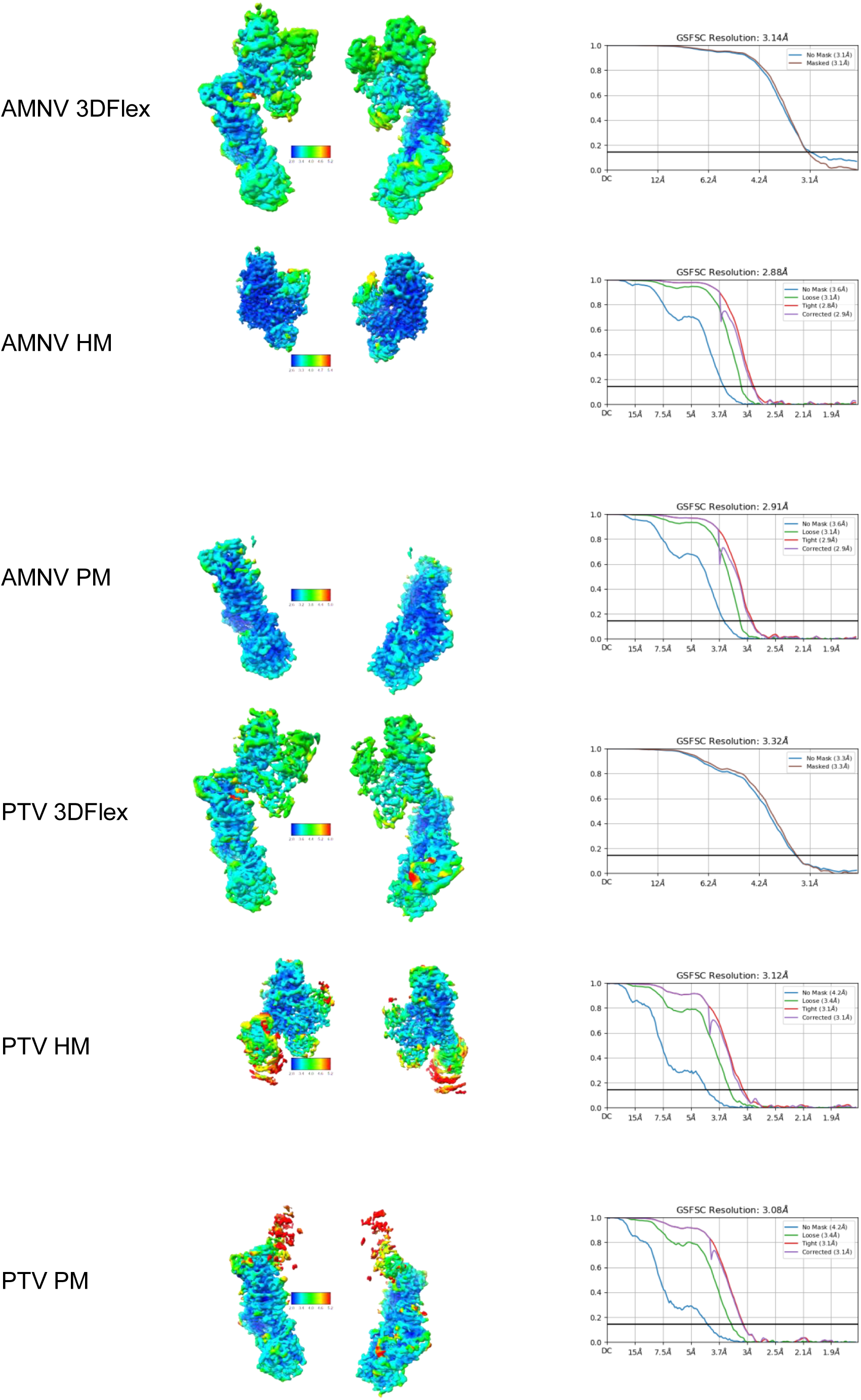
Cryo-EM maps and resolutions of the AMNV and PTV complexes, related to Figure 1.

**Figure S4:**
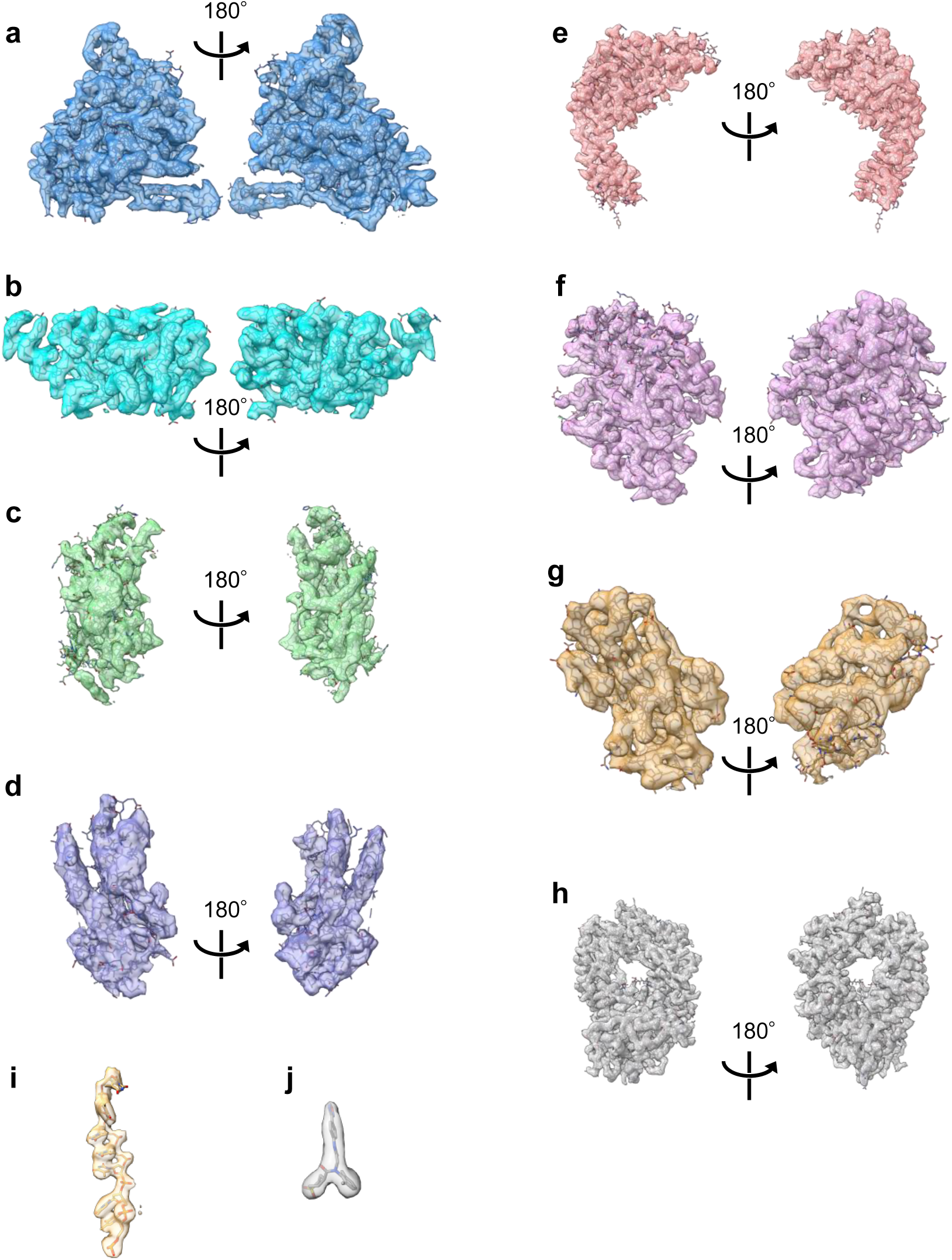
Locally refined maps of individual domains of the AMNV complex, related to Figure 1. **a**, 1A. **b**, 2A. **c**, 2B. **d**, 2C. **e**, SD. **f**, PD. **g**, ZBD. **h**, UL8. **i**, ssDNA. **j**, ANMV.

**Figure S5:**
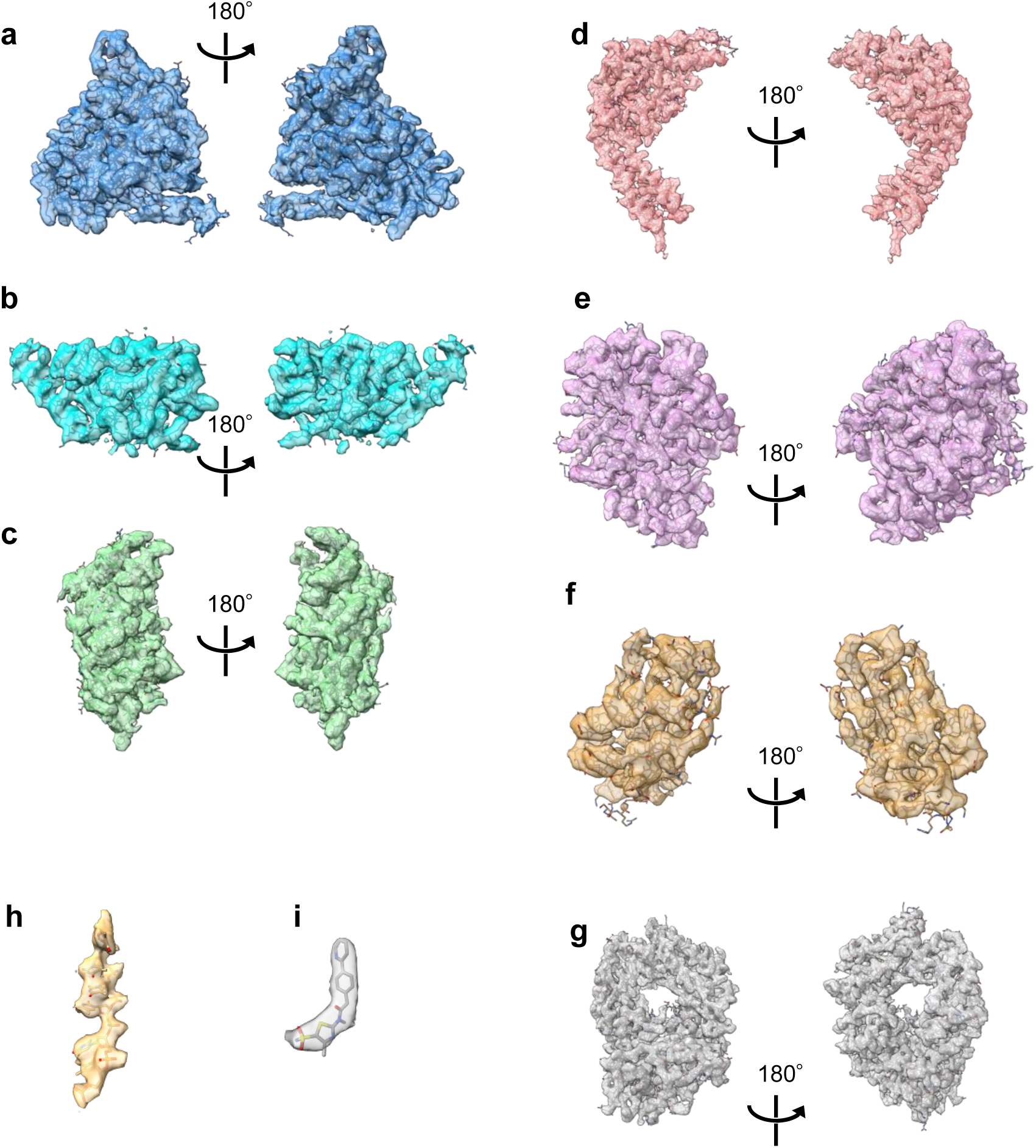
Locally refined maps of individual domains of the PTV complex, related to Figure 1. No model was built for the domain 2C on the locally refined map of the helicase module of the PTV complex because of poor resolution. **a**, 1A. **b**, 2A. **c**, 2B. **d**, SD. **e**, PD. **f**, ZBD. **g**, UL8. **h**, ssDNA. **i**, PTV.

**Figure S6:**
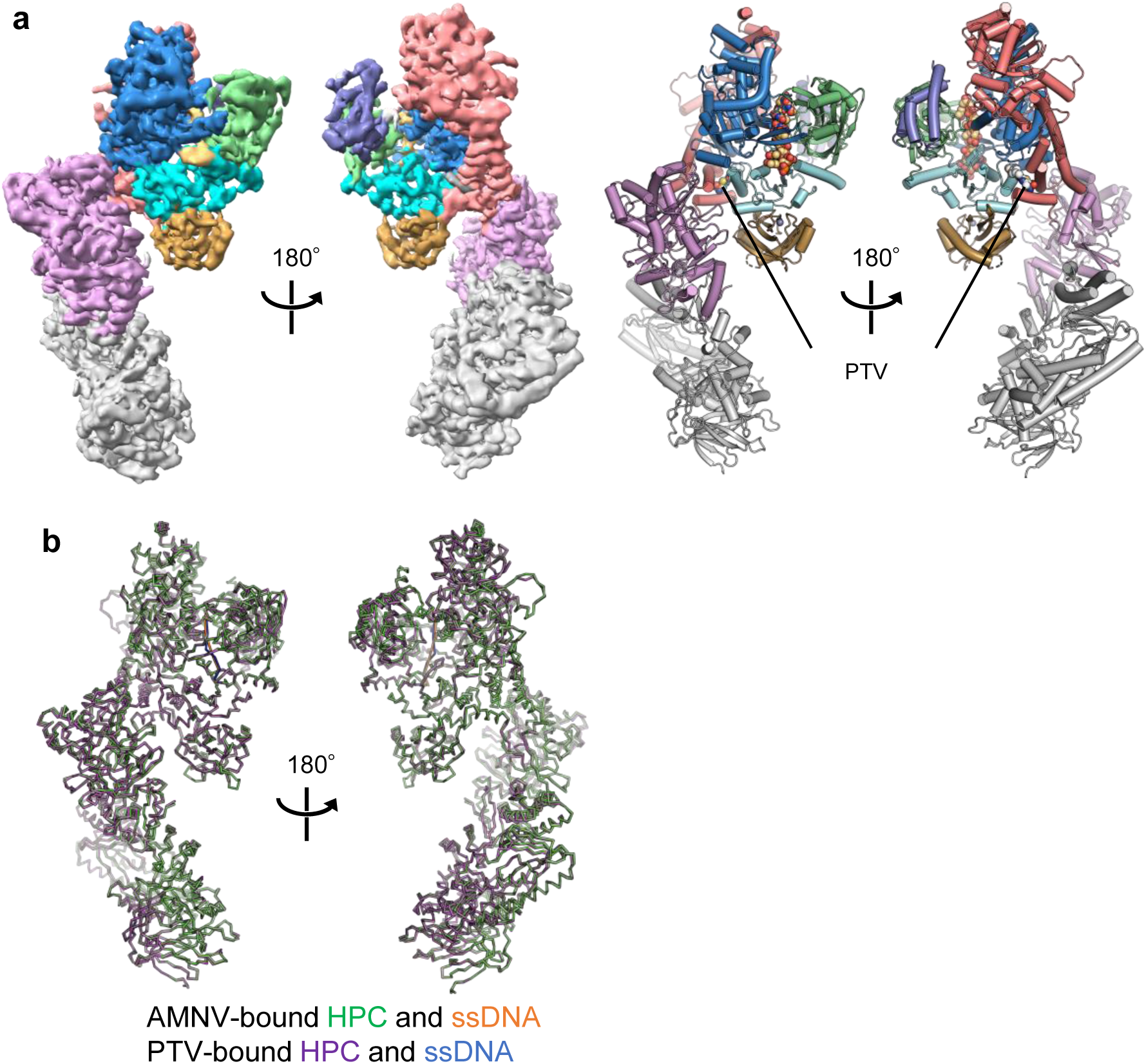
Structure of the PTV-bound HPC, related to Figure 1. **a**, 3DFlex map (calculated using 3DFlex) and the ribbon model. **b**, Structural superposition of the AMNV- and PTV-bound complexes.

**Figure S7:**
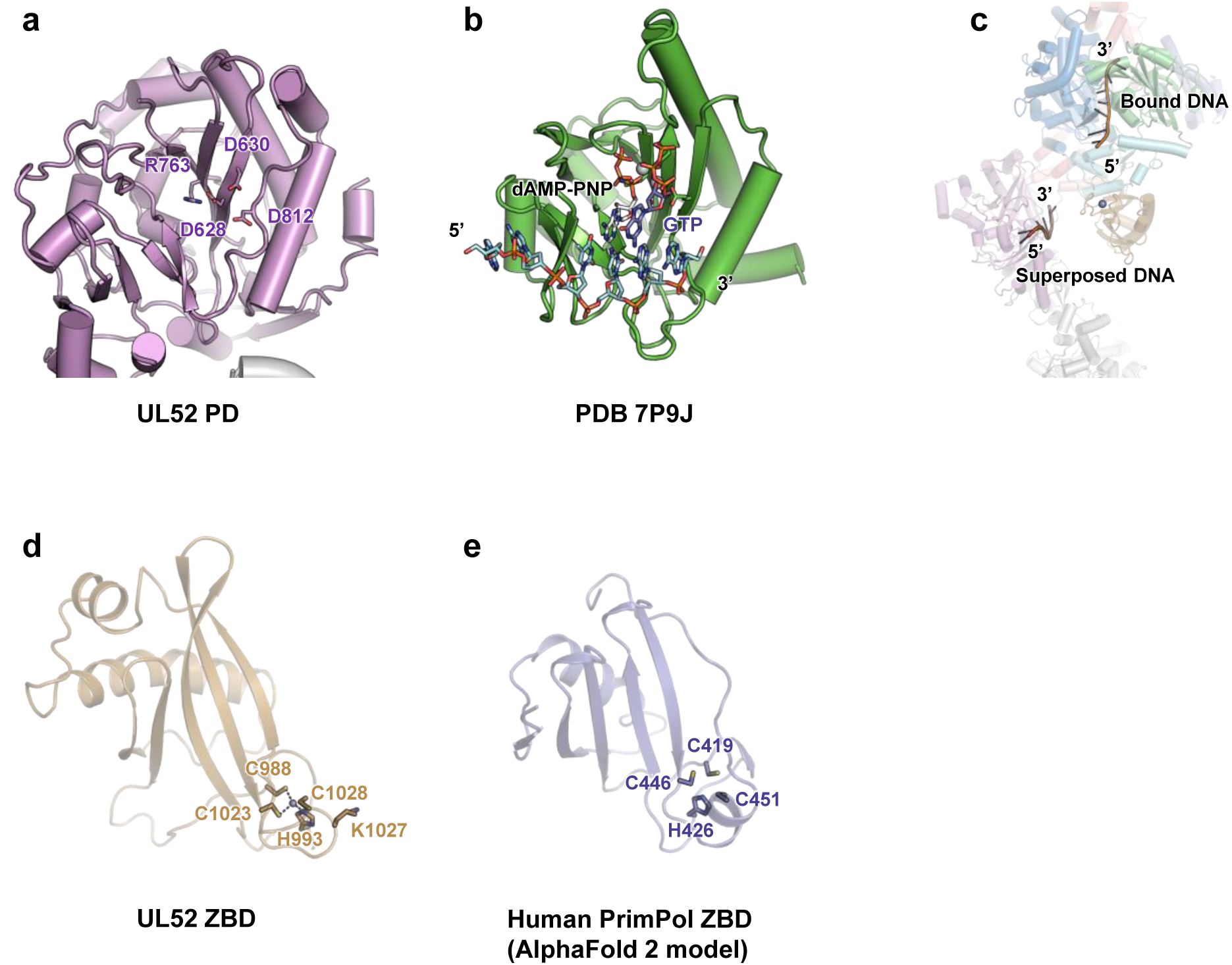
Structures of the PD and ZBD, related to Figure 1. **a**, Structure of the PD. Key catalytic residues conserved among archaeo-eukaryotic primases are shown. **b**, Structure of the primer initiation complex of Prim-Pol domain of CRISPR-associated Prim-Pol (PDB 7P9J). The bound template DNA and two nucleotides (GTP and dAMP-PNP at the initiating and elongating sites, respectively) are also shown. **c**, Superposition of the structures of the AMNV-bound HPC and the template-bound Prime-Pol (only template DNA is shown). **d**, Structure of the ZBD of UL52 in the AMNV-bound HPC. Four zinc-coordinating residues as well as K1027, whose substitution abolishes the activities of HPC, are shown. **e**, A predicted structure of the ZBD (residues 366-468) of human PrimPol (UniProt Q96LW4), downloaded from AlphaFold Protein Structure Database. Possible zinc-coordinating residues are shown.

**Figure S8:**
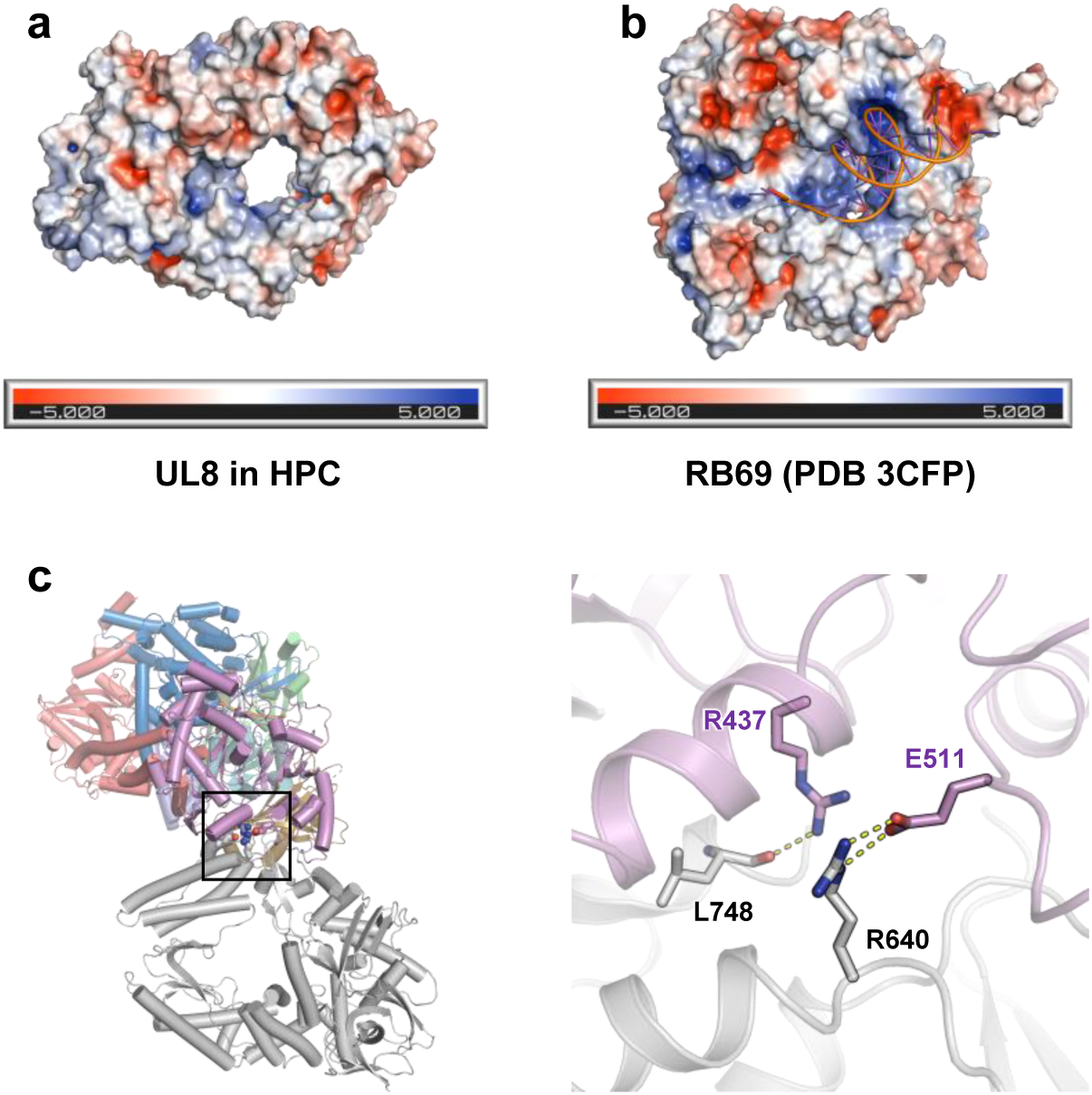
Structure of UL8, related to Figure 1. **a** and **b**, Electrostatic surface potentials of UL8 (**a**) and RB69 (**b**), an α family DNA polymerase, bound with a DNA duplex. **c**, The UL52-UL8 interface. Left, the overall structure of HPC with the UL52-UL8 interface highlighted by a box. Right, A close-up view of the boxed region.

**Figure S9:**
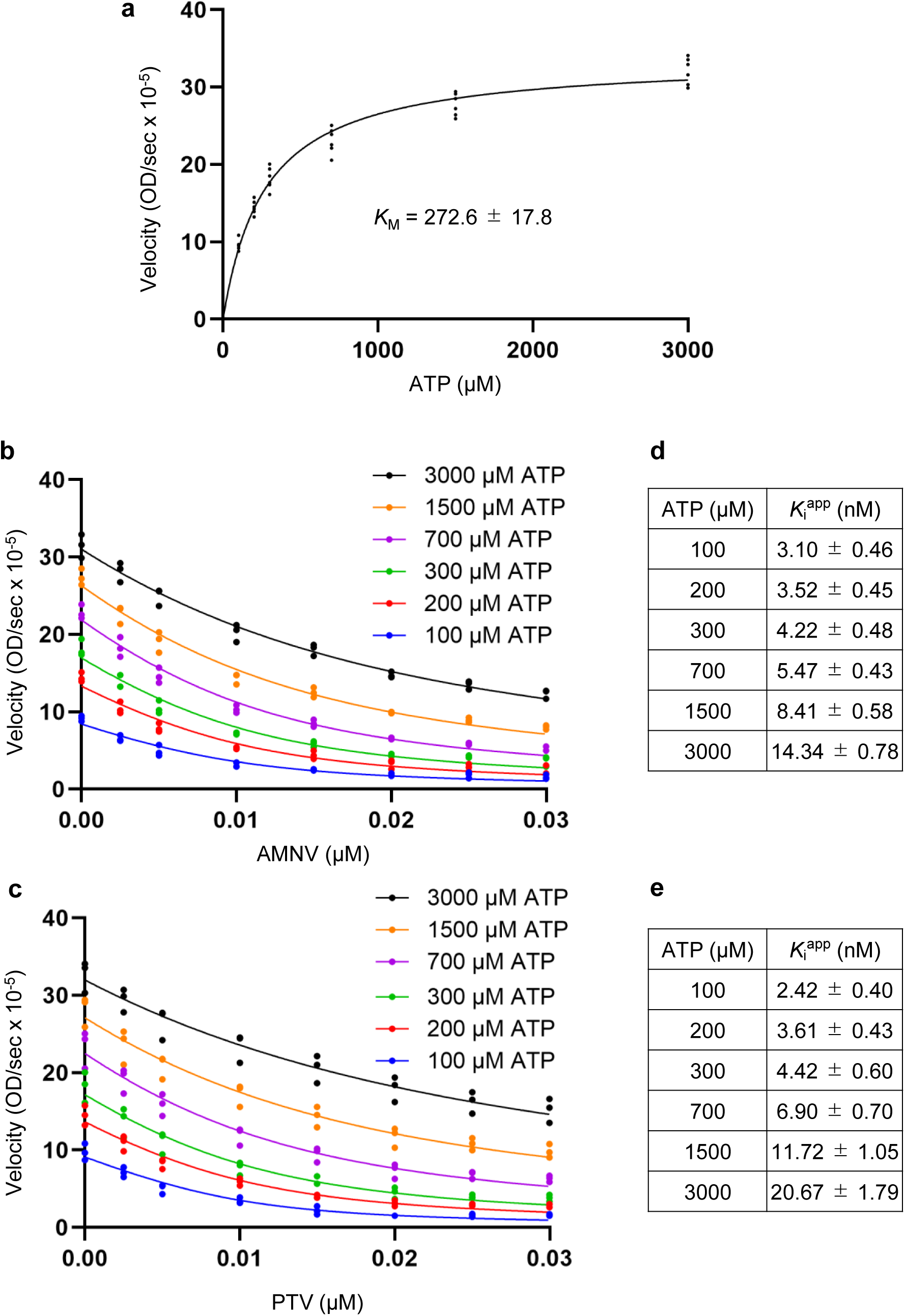
ATPase assays, related to Figure 3. ***a***, Michaelis-Menten plot used to determine the *K_M_* value for ATP from six independent experiments. Standard error of the mean (SEM) is also shown. ***b*** and ***c***, Relationship between reaction velocity and the concentrations of AMNV (***b***) or PTV (***c***) at varying ATP concentrations. ***d*** and ***e***, *K*_i_^app^ values of AMNV (***d***) and PTV € determined under different ATP concentrations. SEM is also shown.

**Figure S10:**
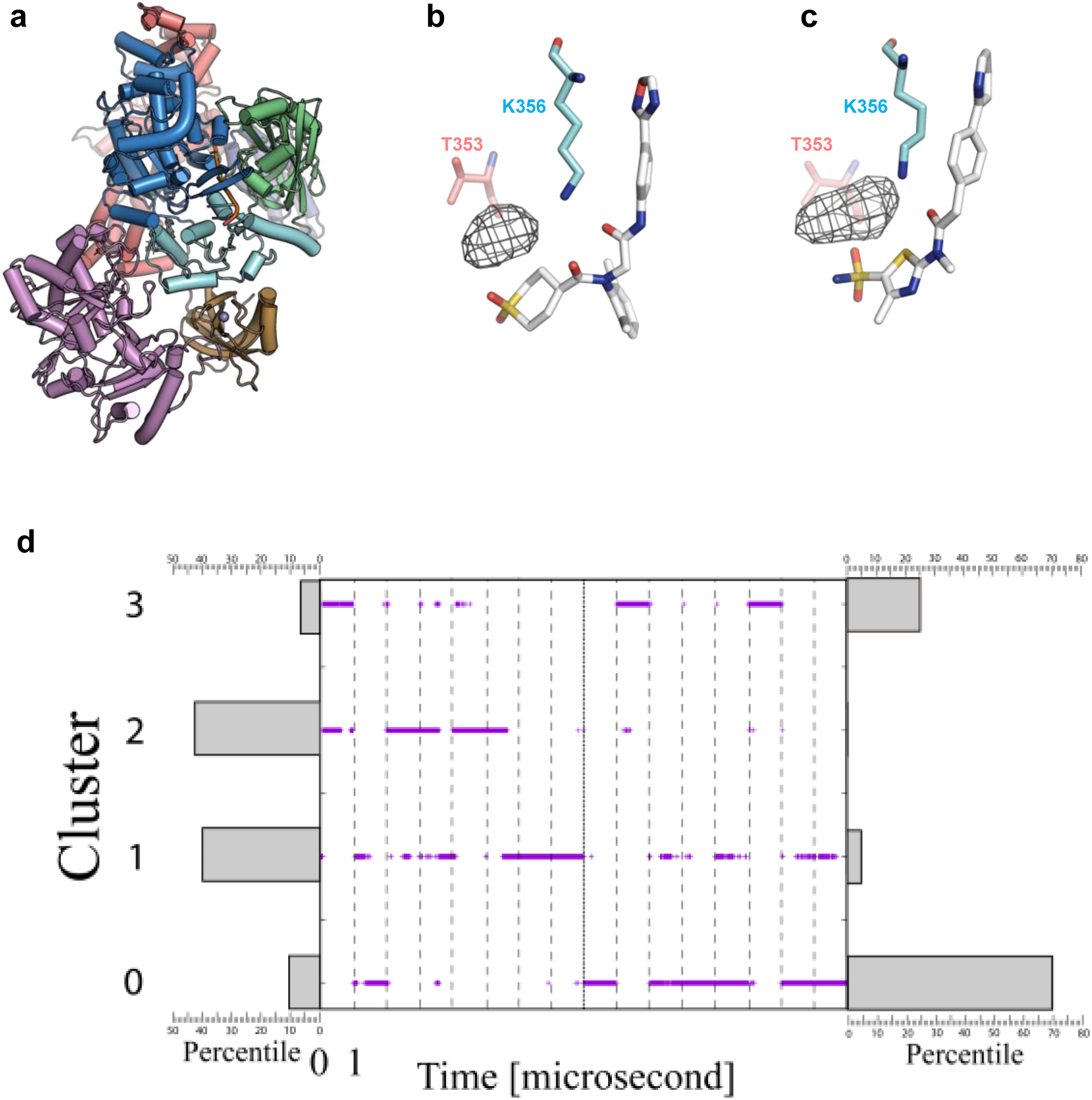
MD analysis, related to Figure 3. a, The modeled structure of the UL5-UL52 subcomplex for MD simulations. b and c, The water density (shown as white mesh, contour level 0.3) within 4.5 Å from the oxygen atoms of the sulfonyl oxygen of AMNV (b) or the sulfonamide NH2 group of PTV (c) are shown. d. The existence percentiles and time distributions of the four clusters for the AMNV and PTV complexes during the MD trajectories are shown on the left and right, respectively. Existence percentiles are represented by bars, while temporal distributions are plotted over time. Vertical dashed lines indicate the one-microsecond MD simulation boundary for each replica (8 replicas per system).

**Figure S11:**
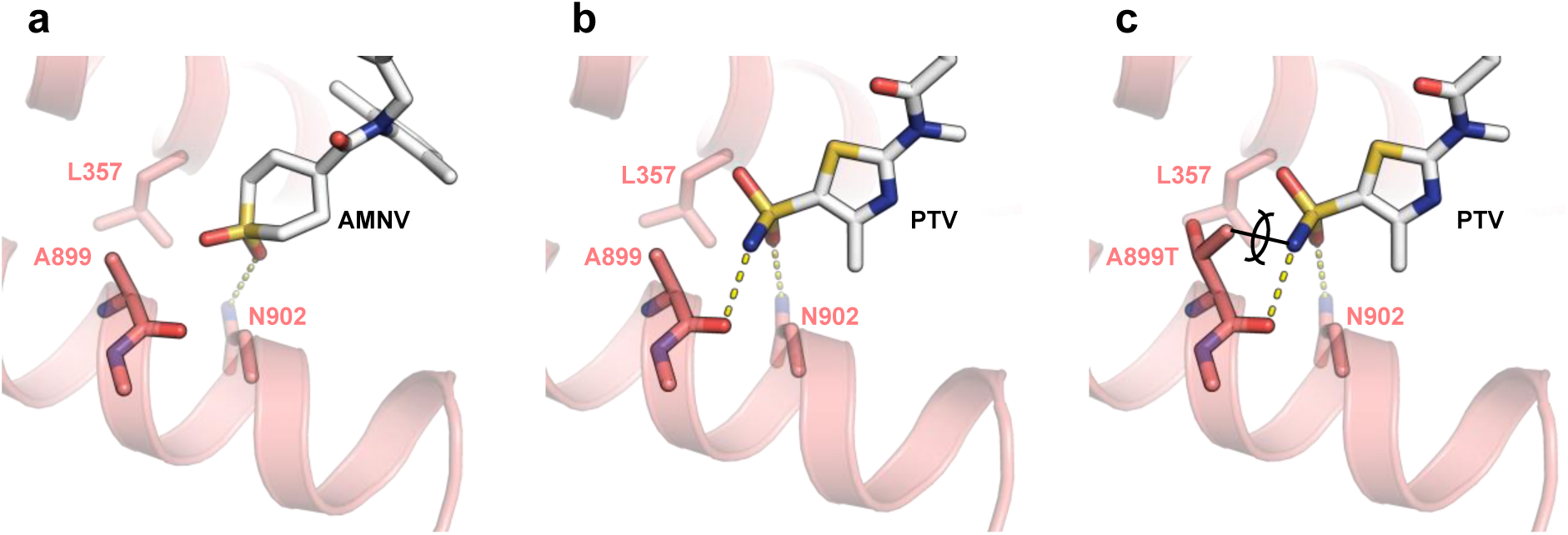
A potential effect of UL52 A899 substitutions on PTV binding, but not AMNV, related to Figure 4. **a**, A hydrogen bond between the sulfonyl group of AMNV and the side chain of N902. **b**, Two hydrogen bonds between the sulfonamide group of PTV and both the side chain of N902 and the carbonyl group of A899. **c**, A structural model of the A899T substitution, which confers resistance to PTV. The Cγ atom of A899T is positioned too close to the amino group of PTV.

**Supplementary Table S1:** Cryo-EM data collection statistics, Related to Figure 1.

|  | AMNV complex | PTV complex |
| --- | --- | --- |
| Magnification | 105,000 | 105,000 |
| Voltage (kV) | 300 | 300 |
| Electron exposure (e<sup>-</sup>/Å<sup>2</sup>) | 49.8 | 50.2 |
| Defocus range (μm) | −0.6 to −1.6 | −0.6 to −1.6 |
| Pixel size (Å) | 0.83 | 0.83 |
| No. of micrographs | 12,466 | 8,102 |

**Supplementary Table S2:** Data processing and refinement statistics.

|  | AMNV<br>3DFlex | AMNV<br>HM | AMNV<br>PM | PTV<br>3DFlex | PTV HM | PTV PM |
| --- | --- | --- | --- | --- | --- | --- |
| **Data processing** |  |  |  |  |  |  |
| EMDB ID | EMD-64478 | EMD-64479 | EMD-64480 | EMD-64481 | EMD-64482 | EMD-64483 |
| Symmetry imposed | $C_1$ | $C_1$ | $C_1$ | $C_1$ | $C_1$ | $C_1$ |
| Particle number | 467,000 | 467,989 | 467,989 | 99,000 | 99,345 | 99,345 |
| Map resolution* | 3.14 | 2.88 | 2.91 | 3.32 | 3.12 | 3.08 |
| **Refinement** |  |  |  |  |  |  |
| PDB ID | 9UT1 | 9UT3 | 9UT4 | 9UT5 | 9UT6 | 9UT7 |
| Non-hydrogen atoms | 18,986 | 10379 | 8607 | 18677 | 9514 | 8607 |
| Protein residues | 2436 | 1298 | 1138 | 2399 | 1195 | 1138 |
| DNA residues | 7 | 7 | 0 | 6 | 6 | 0 |
| Zinc ion | 1 | 1 | 0 | 1 | 1 | 0 |
| Ligand | 1 | 1 | 0 | 1 | 1 | 0 |
| Model vs data CC (mask) | 0.81 | 0.86 | 0.87 | 0.79 | 0.82 | 0.83 |
| Mean B factor ($\text{\AA}^2$) | | | | | | |
| Protein | 174.30 | 142.89 | 116.16 | 155.90 | 130.94 | 117.81 |
| DNA | 118.93 | 84.91 |  | 109.13 | 98.6 |  |
| RMS deviations |  |  |  |  |  |  |
| Bond length ($\text{\AA}$) | 0.003 | 0.003 | 0.003 | 0.002 | 0.003 | 0.002 |
| Bond angles ($^\circ$) | 0.762 | 0.533 | 0.616 | 0.657 | 0.554 | 0.574 |
| Validation |  |  |  |  |  |  |
| MolProbity score | 1.69 | 1.59 | 1.25 | 1.66 | 1.88 | 1.62 |
| Clash score | 6.96 | 7.79 | 4.76 | 5.78 | 9.51 | 8.53 |
| Ramachandran plot |  |  |  |  |  |  |
| Favored (%) | 97.89 | 98.52 | 98.85 | 98.06 | 98.56 | 98.41 |
| Allowed (%) | 2.11 | 1.48 | 1.15 | 1.94 | 1.44 | 1.59 |
| Disallowed (%) | 0 | 0 | 0 | 0 | 0 | 0 |

\*FSC threshold of 0.143 was used.

## Notes

### Competing Interest Statement

The authors have declared no competing interest.

### Summary of Updates

Title updated to focus on the novel insight of the study.

